# Direct binding of arsenicals to nuclear transport factors disrupts nucleocytoplasmic transport

**DOI:** 10.1101/2025.01.13.632748

**Authors:** Emma Lorentzon, Jongmin Lee, Jakub Masaryk, Katharina Keuenhof, Nora Karlsson, Charlotte Galipaud, Rebecca Madsen, Johanna L. Höög, David E. Levin, Markus J. Tamás

## Abstract

Human exposure to arsenicals is associated with devastating diseases such as cancer and neurodegeneration. At the same time, arsenic-based drugs are used as therapeutic agents. The ability of arsenic to directly bind to proteins is correlated with its toxic and therapeutic effects highlighting the importance of elucidating arsenic-protein interactions. In this study, we took a proteomic approach and identified 174 proteins that bind to arsenic in *Saccharomyces cerevisiae*. Proteins involved in nucleocytoplasmic transport were markedly enriched among the arsenic-binding proteins, and we demonstrate that arsenic-binding to nuclear import factors results in their relocation from the nuclear envelope and subsequent aggregation in the cytosol. Similarly, nuclear pore proteins that make up the nuclear pore complex mislocalized and aggregated in arsenic-exposed cells. Consequently, arsenic was shown to inhibit nuclear protein import and export. We propose a model in which arsenic-binding to nuclear transport factors leads to their mislocalization and aggregation, which disrupts nucleocytoplasmic transport and causes arsenic sensitivity.

## INTRODUCTION

Human exposure to the poisonous metalloid arsenic is a global health threat that affects hundreds of millions of people (Chen and Costa, 2021). High concentrations of arsenic in the groundwater have been measured in a large number of countries and long-term exposure is associated with numerous human health problems such as skin disorders, cardiovascular disease, diabetes, cancers of the liver, lung and kidneys, and neurological and neurodegenerative disorders (Chen and Costa, 2021; Rahman et al., 2021; Wysocki et al., 2023). At the same time, arsenic-containing compounds are currently used in anticancer and antiparasitic therapy (Paul et al., 2023).

Pentavalent arsenate, As(V), and trivalent arsenite, As(III) are the most common forms of inorganic arsenic in the environment (Chen and Costa, 2021) and once inside cells, inorganic arsenic can be enzymatically converted into mono-, di-, and trimethylated metabolites (Thomas, 2021). Various forms of arsenic affect cells and living organisms in distinct ways. Due to its chemical similarity to phosphate, As(V) competes with phosphate in biochemical reactions and disrupts adenosine triphosphate (ATP) production. As(III) has high affinity for sulfhydryl groups, such as the thiol groups of cysteine residues, and binding of As(III) and its metabolites to proteins can disrupt protein conformation, function, and interactions (Chen et al., 2019; Kitchin and Wallace, 2008; Shen et al., 2013; Tamás et al., 2014; Vergara-Geronimo et al., 2021; Wysocki et al., 2023). Methylation affects the toxicity of arsenicals as well as their modes of action and protein binding specificities (Shen et al., 2013; Thomas, 2021). For example, As(III) can bind up to three cysteine residues, monomethylarsenite [MAs(III)] can bind two cysteine residues and dimethylarsenite [DMAs(III)] can bind only one cysteine residue (Shen et al., 2013). Arsenic’s ability to bind to proteins is associated with its toxicity, but also with its therapeutic effects. For instance, binding to cysteine residues in the oncoprotein PML-RARα underlies the anticancer activity of arsenic trioxide in patients with acute promyelocytic leukaemia (APL) (Lallemand-Breitenbach et al., 2008; Zhang et al., 2010). Similarly, arsenic-binding to specific kinases and transcriptional regulators is linked to arsenic resistance in yeast and bacteria (Guerra-Moreno et al., 2019; Kumar et al., 2016; Shi et al., 1994). While the toxicity of trivalent arsenite has traditionally been attributed to its interactions with sulfhydryl groups in native (folded) proteins (Kitchin and Wallace, 2008; Shen et al., 2013), recent studies have shown that As(III) also targets non-native proteins impairing their proper folding (Hua et al., 2022; Jacobson et al., 2012; Ramadan et al., 2009; Sapra et al., 2015). In cells, this results in extensive protein misfolding and aggregation which, in turn, has a negative effect on cell proliferation and viability (Andersson et al., 2021; Hua et al., 2022; Ibstedt et al., 2014; Jacobson et al., 2012).

Thus, knowledge of arsenic-protein interactions is key to understand the toxic and therapeutic effects of arsenicals as well as cellular sensing and defence mechanisms. To this end, several large-scale studies have been performed with the aim to identify arsenic-binding proteins (Liu et al., 2023). For example, 360 proteins bound to arsenic *in vitro* using a human proteome microarray (Zhang et al., 2015a) and *in vivo* studies identified 40 arsenic-binding proteins in APL cells (Zhang et al., 2015b), 50 proteins in human breast cancer cells (MCF-7 cell line) (Zhang et al., 2007), 51 proteins in human embryonic kidney epithelial cells (HEK293T) (Dong et al., 2022), and 48 proteins in A549 human lung carcinoma cells (Yan et al., 2016). Follow-up experiments indicated that some of these proteins are *bona fide* targets of arsenic-binding and inhibition (Yan et al., 2016; Zhang et al., 2015a; Zhang et al., 2015b). While identifying the arsenic-binding proteome is a promising approach to address toxicity and resistance mechanisms, the aforementioned *in vivo* studies identified relatively few targets and a comprehensive catalogue of *in vivo* arsenic-protein interactions and the resulting consequences on cell physiology is still lacking.

In this study, we took a proteomic approach and identified 174 proteins that bind to arsenic in budding yeast *Saccharomyces cerevisiae*. Proteins involved in nucleocytoplasmic transport were strongly enriched among the arsenic-binding proteins, and data from follow-up experiments is consistent with a model in which arsenic-binding to nuclear transport factors leads to their mislocalization and aggregation, which disrupts protein transport across the nuclear envelope and causes arsenic sensitivity.

## RESULTS

### Proteome-wide identification of arsenic-binding proteins in yeast

To identify proteins that bind to arsenic *in vivo*, we took an unbiased proteomic approach in *S. cerevisiae* using biotin-conjugated As(III) (hereafter As–biotin) as a probe (Kumar et al., 2016; Lee and Levin, 2019). We previously noted that As-biotin cannot discriminate between As(III)-binding proteins and proteins that bind to MAs(III) due to intracellular conversion of As(III) into MAs(III) (Lee and Levin, 2018; Lee and Levin, 2019). Therefore, we incubated yeast cells that lack the methyltransferase enzyme Mtq2 responsible for As(III) methylation (Lee and Levin, 2018) with 50 µM As–biotin without or with a 10 min pretreatment with 1 mM As(III) or 500 µM MAs(III) as blocking agents (Fig. 1A): the pretreatments were performed to get an indication of which arsenical binds to each protein as binding of As-biotin to a protein is expected to be attenuated in the presence of As(III) or MAs(III), and *mtq2Δ* cells were used to avoid metabolism of the blocking arsenical (Kumar et al., 2016; Lee and Levin, 2018; Lee and Levin, 2019; Lee and Levin, 2022). After cell disruption and As-biotin pull-down with streptavidin-agarose beads, candidate arsenic-binding proteins were eluted, separated by SDS-PAGE electrophoresis, and identified by microcapillary liquid chromatograph-tandem mass spectrometry (LC/MS/MS) (Fig. 1A). As a control, we performed pull-downs using cells that had not been incubated with As-biotin. In total, 776 proteins were identified in at least one of the conditions (Supplementary Table S1). To select candidate arsenic-binding proteins, we filtered the 776 proteins using the following criteria: (1) no peptide present in the control, (2) ≥5 unique peptides identified after As-biotin pull-down, and (3) ≥2-fold reduction of signal/peptide intensity when competitor As(III) or MAs(III) was present during pull-down. Applying these stringent filtering criteria gave a list of 174 candidate arsenic-binding proteins (Table S2). To our knowledge, this represents the largest set of *in vivo* arsenic-binding proteins reported to date.

**Figure 1.**
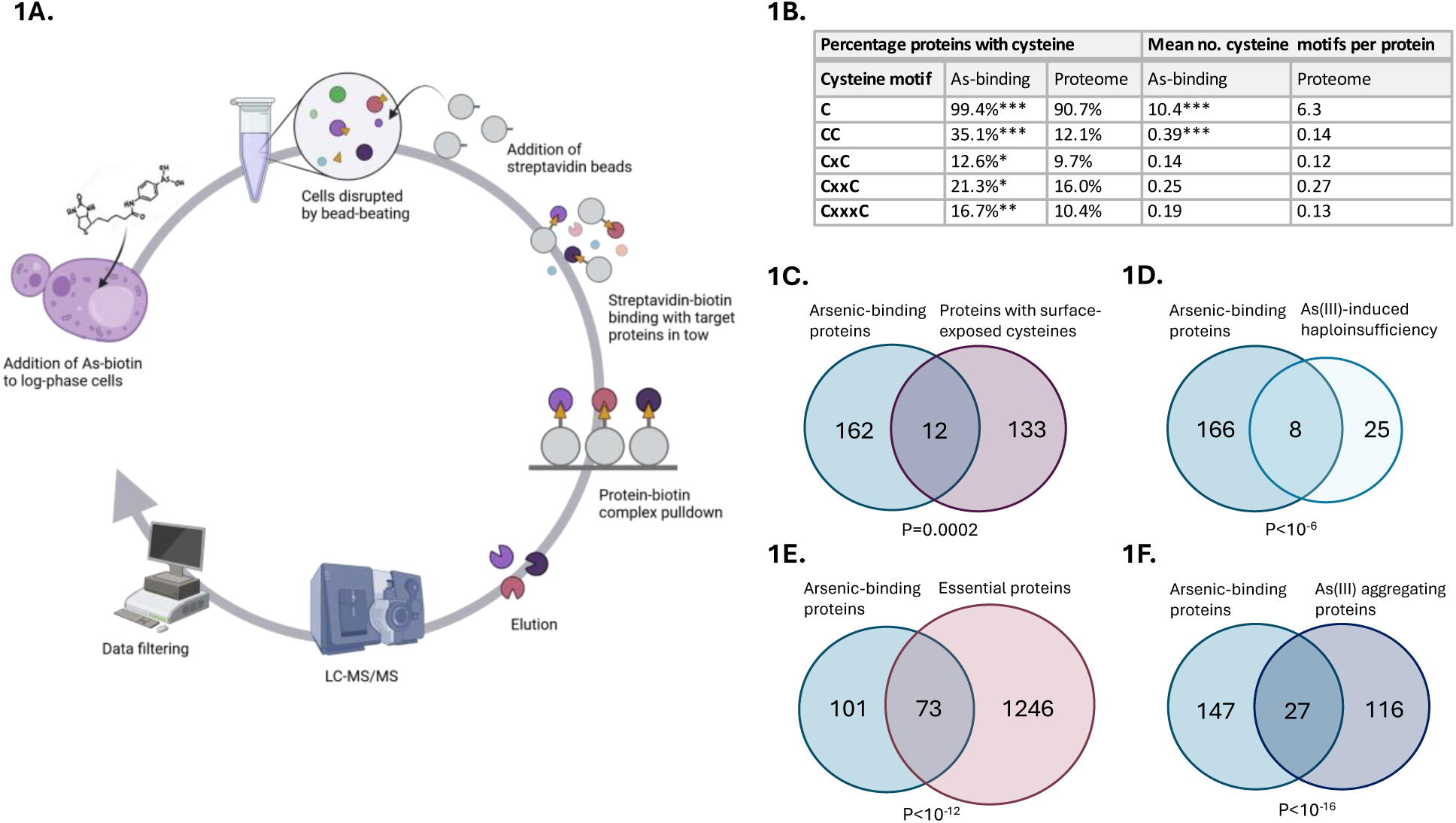
Proteome-wide screen identifies arsenic-binding proteins and toxicity targets. **1A**. Workflow. Yeast cells (*mtq2Δ*) were incubated with 50 µM As-biotin without or with a 10 min pretreatment with 1 mM As(III) or 500 µM MAs(III) as blocking agents. After cell disruption and protein pull-down using streptavidin beads, the proteins present in the pull-down were identified using LC-MS/MS. The data was filtered using the following stringent criteria: (1) no peptide present in the control, (2) ≥5 unique peptides identified per protein after As-biotin pull-down, and (3) ≥2-fold reduction of signal/peptide intensity when competitor As(III) or MAs(III) was present during pull-down. A total of 174 candidate arsenic-binding proteins were identified. **1B**. Cysteine content and motifs in arsenic-binding proteins versus a proteome of around 5800 proteins (Ho et al., 2018). x represents any amino acid present between the cysteine residues in a motif. Significance was calculated by the hyper-geometric test, and *P*-values are according to: * < 0.05, ** < 0.01, and *** < 0.001. **1C-F**. Venn diagrams show the overlap between arsenic-binding proteins and: **C)** proteins containing surface-exposed cysteine residues (Marino et al., 2010), **D)** As(III)-sensitive heterozygous diploid knockout mutants (Pan et al., 2010), **E)** essential proteins in *S. cerevisiae* (extracted from SGD (Wong et al., 2023)), and **F)** proteins aggregating during As(III) exposure (Ibstedt et al., 2014; Jacobson et al., 2012). The significance of the overlaps between the datasets was calculated by the hyper-geometric test and the corresponding *P*-values are indicated.

Several of the 174 yeast proteins have human orthologues that were previously reported to bind to arsenic in large-scale *in vitro* and *in vivo* screens (Dong et al., 2022; Zhang et al., 2015a; Zhang et al., 2007) including subunits of the chaperonin TRiC/CCT complex involved in protein folding, metabolic enzymes such as glycerol-3-phosphate dehydrogenase, aldehyde dehydrogenase and members of the pyruvate dehydrogenase complex, proteins involved in DNA replication including components of the minichromosome maintenance (MCM) complex, α- and β-tubulin, and ribonucleotide reductase implicated in DNA synthesis and repair. Thus, these proteins may represent evolutionarily conserved arsenic-binding targets. Of these, the chaperonin TRiC/CCT (Pan et al., 2010), tubulin (Zhang et al., 2007), and pyruvate dehydrogenase (Bergquist et al., 2009; Peters et al., 1946) have been proposed to be direct toxicity targets.

The majority of the 174 proteins (103 proteins, 59%) had ≥2-fold reduced signal/peptide intensity in the presence of As(III) as well as MAs(III), suggesting that they may bind both arsenicals (Table S2). 48 proteins (28%) reached the threshold of ≥2-fold reduction in signal/peptide intensity only in presence of MAs(III) while 23 (13%) reached the threshold only in presence of As(III), suggesting that these proteins preferentially bind to either MAs(III) or As(III), respectively. As(III) and MAs(III) preferentially bind to the thiol group of cysteine residues in proteins (Kitchin and Wallace, 2008; Shen et al., 2013; Zhang et al., 2015a), and virtually all 174 proteins (99.4%) contained at least one cysteine compared to 90.7% in the yeast proteome (*P*<10^-7^) (Fig. 1B). The arsenic-binding set was also significantly enriched for proteins with cysteines adjacent or proximal to other cysteines (CC, CxC, CxxC and CxxxC motifs), and the mean number of cysteines and CC motifs per protein was significantly higher in arsenic-binding proteins compared to the proteome (Fig. 1B). Additionally, we observed a significant overlap between the arsenic-binding proteins and a set of 145 yeast proteins that possess surface-exposed reactive cysteines (12 proteins, *P*=0.0002) (Marino et al., 2010) (Fig. 1C). As(III) has been shown to bind to proteins containing zinc finger motifs, specifically to C3H1 and C4 motifs (Vergara-Geronimo et al., 2021; Zhou et al., 2011). 12 of the 174 proteins (*P*=0.11) in our dataset are putative zinc-binding proteins of which 8 are predicted to contain C3H1 and C4 motifs (Wang et al., 2018). In sum, the As-biotin probe identified proteins that bind to As(III) and MAs(III) or both arsenicals, and our findings reinforce the strong preference of As(III)/MAs(III) for cysteine residues in proteins *in vivo*.

### Protein binding as a possible toxicity mechanism

It has been postulated that trivalent arsenic causes toxicity via protein binding, inactivating or depleting important cellular functions (Chen et al., 2019; Kitchin and Wallace, 2008; Shen et al., 2013; Tamás et al., 2014; Vergara-Geronimo et al., 2021; Wysocki et al., 2023). However, only few direct toxicity targets and mechanisms have been described to date. One way to pinpoint candidate toxicity targets is to perform drug-induced haploinsufficiency profiling (HIP) assays (Giaever et al., 1999; Lum et al., 2004). A previous yeast HIP study identified 33 As(III) sensitive heterozygous diploid knockout mutants (Pan et al., 2010) of which 8 encode proteins that were present in the arsenic-binding set (*P*<10^-6^) (Fig. 1D) including components of TRiC/CCT (Cct1, Cct4, Cct5, Cct7), α-tubulin (Tub3), the nuclear pore protein Nup145, adenine-requiring Ade12, and the serine palmitoyltransferase Lcb1. These proteins may represent *bona fide* arsenic toxicity targets, as proposed for TRiC/CCT (Pan et al., 2010) and tubulin (Zhang et al., 2007).

Another way to identify direct toxicity targets is to integrate chemical-genetic and genetic interaction data (Parsons et al., 2004). For this, we retrieved negative genetic interactors of selected arsenic-binding proteins and asked whether the sets of negative genetic interactors are enriched for As(III) sensitive mutants. Indeed, we observed significant enrichments in As(III) sensitivity among negative genetic interactors of selected arsenic-binding protein-encoding genes involved in transport across the nuclear envelope (*KAP121/PSE1* (*P*=0.0004), *NUP84* (*P*<10^-20^)), components of TRiC/CCT (*CCT1* (*P*<10^-13^), *CCT5* (*P*<10^-7^)), the alpha subunit of pyruvate dehydrogenase *PDA1* (*P*<10^-13^), the translation regulator *GCN20* (*P*=0.004), and the serine palmitoyltransferase *LCB1* (*P*=0.0002) (Fig. S1). Thus, the tested arsenic-binding proteins might represent *bona fide* toxicity targets, as proposed for TRiC/CCT (Pan et al., 2010) and pyruvate dehydrogenase (Bergquist et al., 2009; Peters et al., 1946).

We noted that a substantial fraction of the arsenic-binding proteins is essential for cell viability (73 proteins, 42%, *P*<10^-12^) (Fig. 1E), suggesting that arsenic-binding might drain the active pool of these essential proteins, resulting in poor growth or survival of yeast cells during arsenic stress. The set of essential arsenic-binding proteins included proposed toxicity targets such as TRiC/CCT (Pan et al., 2010) and tubulin (Zhang et al., 2007) and novel candidate targets such as proteins involved in nucleocytoplasmic transport (Fig. S1; see further).

One way arsenic binding affects protein function is by interfering with their folding, thereby preventing proteins from reaching their native fold and hence, their active conformation (Andersson et al., 2021; Hua et al., 2022; Ibstedt et al., 2014; Jacobson et al., 2012). We found a significant overlap between the set of arsenic-binding proteins and a set of 143 proteins that aggregated in As(III)-exposed yeast cells (27 proteins, *P*<10^-16^) (Ibstedt et al., 2014; Jacobson et al., 2012) (Fig. 1F), suggesting that arsenic-binding to these proteins results in their misfolding and aggregation.

In sum, our findings suggest that a large fraction of the 174 arsenic-binding proteins identified here may represent *bona fide* toxicity targets and support the notion that protein binding is a major arsenic toxicity mechanism, by affecting protein folding and/or activity. Thus, integrating arsenic-protein binding data with other datasets is a powerful approach to identify novel toxicity targets.

### Arsenic binds to proteins involved in nucleocytoplasmic transport

We next addressed if specific categories of protein functions are overrepresented among the arsenic-binding proteins. Gene ontology (GO) analysis revealed that this set was enriched in processes associated with protein import into the nucleus, chaperonin-mediated protein folding, carboxylic acid metabolic processes, nuclear pore localization and organization, DNA unwinding involved in DNA replication, tRNA transport, methylation and aminoacylation, and sphingosine and long-chain fatty acid metabolism (Fig. 2A). Remarkably, the set of arsenic-binding proteins was markedly enriched for functions in nucleocytoplasmic transport and included 8 out of the 11 importins (Kap104, Kap114, Kap122, Kap123, Kap121/Pse1, Sxm1/Kap108, Nmd5/Kap119, Mtr10/Kap111) present in *S. cerevisiae*, several exportins (Cse1/Kap109, Crm1/Kap124, Msn5/Kap142), and numerous nuclear pore proteins (Nup84, Nup85, Nup120, Nup133, Nup145, Nup157, Nup170, Nup188, Nup192) (Table S2). This strong enrichment together with our genetic analyses above implicating some of these proteins as direct toxicity targets (Nup145, Nup84, Kap121/Pse1), raised the possibility that nucleocytoplasmic transport is a key target of arsenic in living cells.

**Figure 2.**
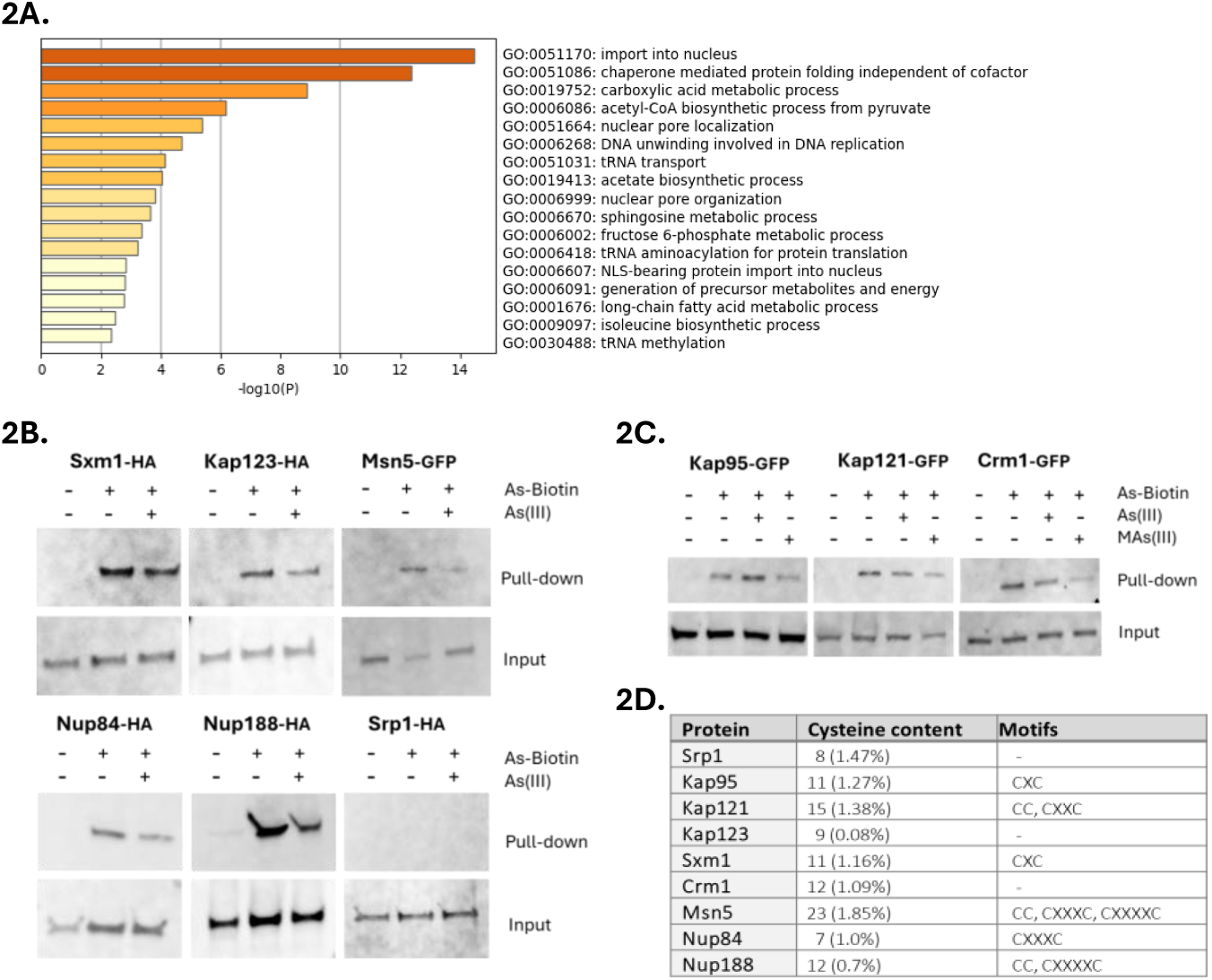
Arsenic binds to proteins involved in nucleocytoplasmic transport. **2A**. Bar plots of overrepresented GO-terms in the arsenic-binding protein set using Metascape (Zhou et al., 2019). **2B**. Cells expressing HA-tagged or GFP-tagged versions of Sxm1, Kap123, Msn5, Nup84, Nup188 and Srp1 were incubated with 50 µM As-biotin followed by cell disruption and protein pull-down using streptavidin beads. The proteins were detected by Western blotting using anti-HA and anti-GFP antibodies. Cells were pretreated with 1 mM As(III) as blocking agent as indicated. The loading control (Input) represents the total lysate. The blots shown are representative of at least two biological repeats. **2C**. As-biotin pulldown assays were performed as in 2B using cells expressing GFP-tagged Kap95, Kap121/Pse1 and Crm1. The proteins were detected by Western blotting using an anti-GFP antibody. Cels were pretreated with 1 mM As(III) or 1 mM MAs(III) as blocking agents as indicated. The blots shown are representative of at least two biological repeats. **2D**. Cysteine content and motifs in the listed proteins. x represents any amino acid present between the cysteine residues in a motif.

Importins are nuclear transport factors that bind to cargo proteins containing nuclear localization signals (NLS) in the cytoplasm and facilitate their passage through the nuclear pore complex (NPC) into the nucleus, exportins mediate the export of cargo proteins back to the cytoplasm, while nuclear pore proteins (Nups) constitute the NPC (Aitchison and Rout, 2012; Wing et al., 2022). Importins and exportins belong to the karyopherin (Kap) family of nuclear transport factors. While most karyopherins can bind to their cargos directly (karyopherin β), in the case of the heterodimeric α/β complex, it is karyopherin α that binds to the cargo with karyopherin β stabilizing/enhancing this interaction (Aitchison and Rout, 2012; Wing et al., 2022). To validate arsenic-binding to selected importins (karyopherin βs Kap123, Kap121/Pse1, Sxm1/Kap108), exportins (karyopherin βs Crm1/Kap124, Msn5/Kap142) and Nups (Nup84, Nup188), we either introduced plasmids that expressed HA-tagged versions of the corresponding genes into yeast cells or used cells that harboured GFP-tagged versions of the genes in their genomes, and performed As-biotin pulldown assays. All tested proteins bound to As-biotin and this binding was attenuated to various degrees in the presence of competitor As(III) or MAs(III) (Figs. 2B, 2C). Thus, these proteins directly bind arsenic *in vivo* in form of As(III) and/or MAs(III). We also chose to test arsenic-binding to the karyopherin α Srp1/Kap60 and the karyopherin β Kap95 even though they were not present in the hit list (no peptides were found for Srp1/Kap60 while Kap95 was below the threshold), because these proteins constitute the heterodimeric α/β complex that plays a key role in nuclear transport of NLS-containing proteins in yeast (Aitchison and Rout, 2012). Kap95 readily bound to As-biotin and this binding was unaffected by competitor As(III) but clearly attenuated in the presence of MAs(III) (Fig. 2C). Thus, Kap95 binds to arsenic in form of MAs(III) *in vivo*. In contrast, Srp1 did not bind to As-biotin (Fig. 2B).

Most of the tested proteins (Sxm1, Kap95, Kap121/Pse1, Msn5, Nup84, Nup188) have adjacent or proximal cysteines in their primary sequence (Fig. 2D). Analyses of their known or predicted 3D structures revealed that most of these proteins contain cysteine pairs within approximately 5Å of each other (Fig. S2), making them suitable substrates for As(III)/MAs(III) in their native folded structures. In contrast, Kap123 and Crm1 lack adjacent or proximal cysteines in their primary sequence, raising the question of how As(III) and/or MAs(III) bind to these proteins. Inspection of their 3D structures revealed the presence of proximal cysteines that could potentially serve as binding sites in both proteins (Fig. S2). Finally, Srp1/Kap60 does not have cysteine motifs in its primary sequence and the closest cysteines in its 3D structure are separated by approximately 10Å with the thiol groups pointing in opposite directions (Fig. S2), explaining why this protein is a poor substrate for As(III)/MAs(III).

### Importins mislocalize and aggregate in As(III)-exposed cells

Having established that arsenic binds to individual importins, we next addressed the consequence(s) of this binding. First, we monitored the localization of chromosomally integrated Kap95-GFP. We chose to focus on Kap95 because: (1) it plays a central role in nuclear transport of NLS-containing proteins in yeast (Aitchison and Rout, 2012), (2) integration of chemical-genetic and genetic interaction data suggested that Kap95 may be a direct arsenic toxicity target (Fig. S1), and (3) the human orthologue of Kap95, Importin 90/KPNB1, was identified as a candidate arsenic-binding protein in MCF-7 cells (Zhang et al., 2007). In untreated (control) cells, chromosomally integrated Kap95-GFP localized around the yeast nuclear envelope (NE) (Fig. 3A, 3B). In As(III)-exposed cells, NE localization of Kap95-GFP was disrupted and the protein was instead found in distinct foci that were dispersed throughout the cytosol in the majority of cells (Figs. 3A, 3B). Cytosolic Kap95-GFP foci were also formed when the translation inhibitor cycloheximide (CHX) was added at the same time as As(III) (Fig. 3B), suggesting that As(III) impacts the localization of the native (folded) form of Kap95. Kap95 forms a heterodimeric nuclear transport receptor together with Srp1 (Aitchison and Rout, 2012). As for Kap95, As(III) stress altered the distribution of chromosomally integrated Srp1-GFP from the NE to distinct cytosolic foci (Fig. 3A) both in the absence and presence of CHX (Fig. S3A), which is consistent with the two proteins forming a heterodimer.

**Figure 3.**
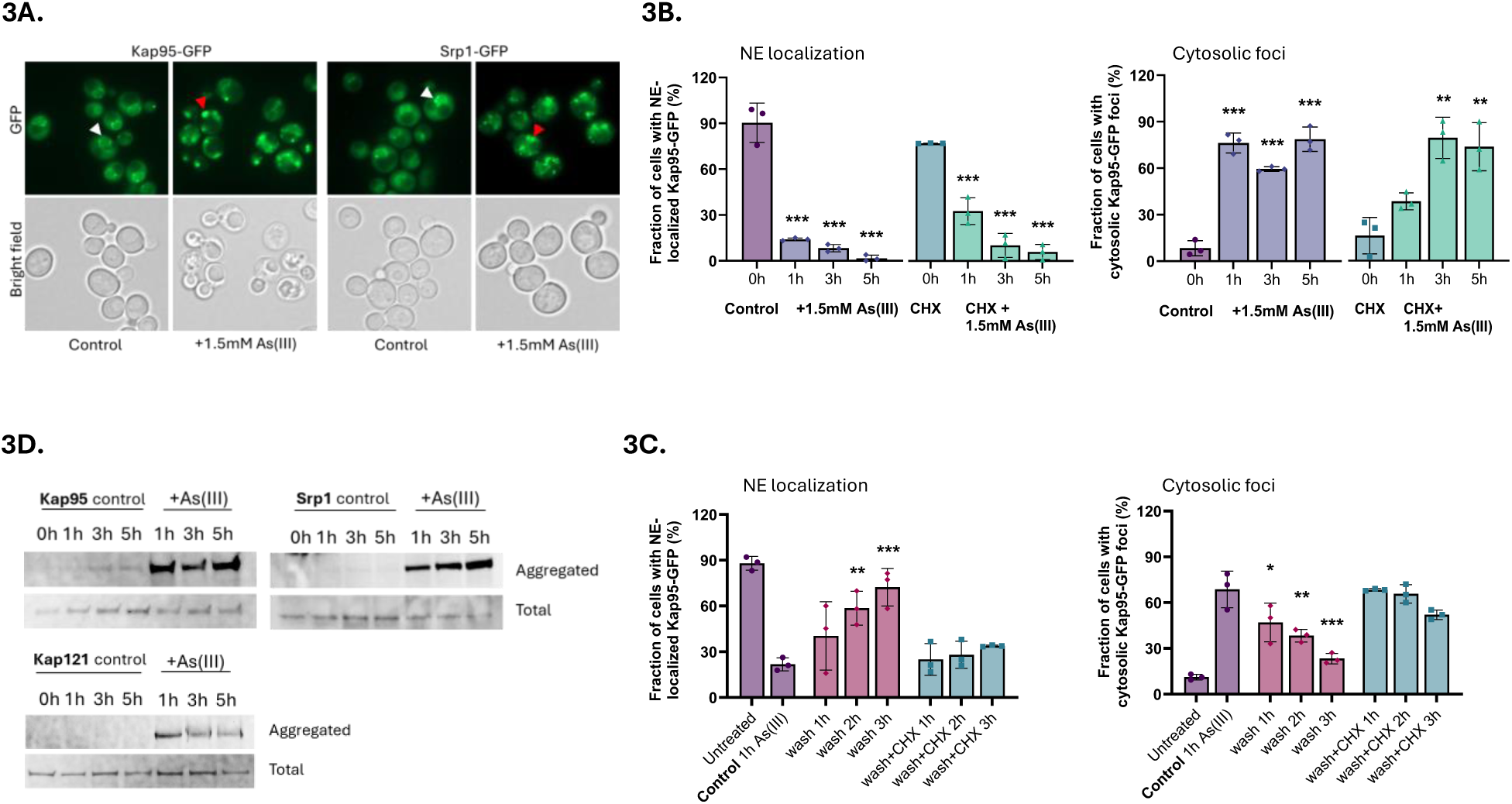
Importins mislocalize and aggregate in As(III)-exposed cells. **3A**. Localization of GFP-tagged Kap95 and Srp1. In unexposed (control) cells the proteins are located around the NE (white arrowheads). After As(III) exposure (1.5 mM) for 1 h, the proteins are found in distinct foci dispersed throughout the cytosol (red arrowheads). Images shown are representative of three biological repeats of 100 cells each. **3B**. Quantification of Kap95-GFP NE localization (left panel) and cytosolic foci formation (right panel) in the absence and presence of 1.5 mM As(III) and/or 0.2 mg/ml cycloheximide (CHX). Kap95–GFP distribution was scored by fluorescence microscopy and quantified by visual inspection. The bars represent the mean ± standard deviation (SD) of three independent biological repeats of a total of 300 cells. Significance was calculated using un-paired two-tailed student’s t-test with either the untreated control (for just As(III) exposure) or CHX (for CHX+As(III)-treated cells) as the comparison. *P*-values are according to: ** > 0.01, *** > 0.001. **3C**. Cells were exposed to 1.5 mM As(III) for 1 h, then washed twice and resuspended in medium without As(III) in the presence or absence of 0.2 mg/ml CHX. Kap95–GFP distribution was scored and quantified as in 3B. The bars represent the mean ± SD of three independent biological repeats of a total of 300 cells. Significance was calculated using un-paired two-tailed student’s t-test of three independent biological replicates, with 1 h As(III) exposed cells as the control. *P*-values are according to: * > 0.05, ** > 0.01, *** > 0.001. **3D**. Kap95, Srp1, and Kap121/Pse1 aggregate in the presence of As(III). Cells expressing GFP-tagged proteins were left untreated (control) or exposed to 1.5 mM As(III), lysed, and the total and aggregated protein fractions were isolated followed by Western blot analysis using an anti-GFP antibody. The blots shown are representative of at least two biological repeats.

Controlled formation of protein condensates can be used by cells for various physiological purposes whereas aggregation of misfolded proteins represents an irreversible loss of protein function (Alberti and Hyman, 2021). To address whether formation of cytosolic Kap95-GFP and Srp1-GFP foci are reversible, we monitored their localization after As(III) exposure during 1 h followed by As(III) washout. NE localization of both proteins slowly recovered after As(III) washout, and this recovery largely coincided with the disappearance of cytosolic Kap95-GFP (Fig. 3C) and Srp1-GFP (Fig. S3B) foci. CHX, added after the washing step, slowed down or prevented the recovery of Kap95-GFP and Srp1-GFP at the NE as well as the disappearance of cytosolic Kap95-GFP and Srp1-GFP foci (Figs. 3C, S3B), indicating that foci reversal and signal recovery at the NE require *de novo* protein synthesis. Thus, formation of Kap95-GFP and Srp1-GFP foci may not be a regulated process that cells use to quickly recover once As(III) stress is relieved.

Next, we addressed whether the Kap95-GFP and Srp1-GFP foci represent aggregated forms of these proteins. For this, we isolated total and aggregated proteins by differential centrifugation and separated the proteins in each fraction by SDS-PAGE followed by immunoblotting with an anti-GFP antibody. Both Kap95-GFP and Srp1-GFP were present in the aggregated protein fractions isolated from As(III)-exposed cells while these proteins were largely absent in the aggregated protein fractions of unexposed cells (Fig. 3D). This finding indicates that Kap95 and Srp1 aggregate in the presence of As(III) and that the cytosolic foci likely represent aggregated forms of these proteins. The data also suggest that arsenic-binding to Kap95, in form of MAs(III), is sufficient to induce mislocalization and aggregation of both Srp1 and Kap95 in the heterodimeric α/β complex. Importantly, the karyopherin βs Kap121/Pse1 (Fig. 3D) and Kap123 (Jacobson et al., 2012) also aggregated during As(III) exposure. We conclude that arsenic-binding to nuclear import receptors leads to their relocation from the nuclear envelope and subsequent aggregation in the cytosol.

### As(III) affects Nup localization, NE morphology, and NPC numbers

The arsenic-binding Nups identified in this study are located in the outer ring (Nup84, Nup85, Nup120, Nup133, Nup145) and the inner ring (Nup157, Nup170, Nup188, Nup192) of the NPC (Fig. 4A). Nup145 is, after proteolytic cleavage, present in the outer ring (Nup145C fragment) and in the NPC core (Nup145N) as one of several so-called FG-Nups that are directly responsible for nucleocytoplasmic transport (Aitchison and Rout, 2012; Wing et al., 2022). Similar to the tested importins, As(III) affected the localization of chromosomally integrated Nup84-GFP and Nup188-GFP. In untreated (control) cells, Nup84-GFP was localized in patches in the NE (Fig. 4B). During As(III) exposure, Nup84-GFP was visible as cytosolic foci and the fluorescence signal appeared to extend from the NE into the cytosol in a substantial fraction of the cells (Figs. 4B, 4C). Likewise, Nup188-GFP was visible in a punctuate pattern in the NE in unexposed cells whereas cytosolic foci and fluorescence signal extensions from the NE into the cytosol were observed in As(III)-exposed cells (Figs. 4B, 4C). CHX did not prevent As(III)-induced Nup84-GFP foci formation (Fig. S4A), suggesting that As(III) affects the localization of the native (folded) form of Nup84. These findings imply that arsenic-binding to Nup84 and Nup188 leads to their mislocalization and that arsenic may cause NE deformations.

**Figure 4.**
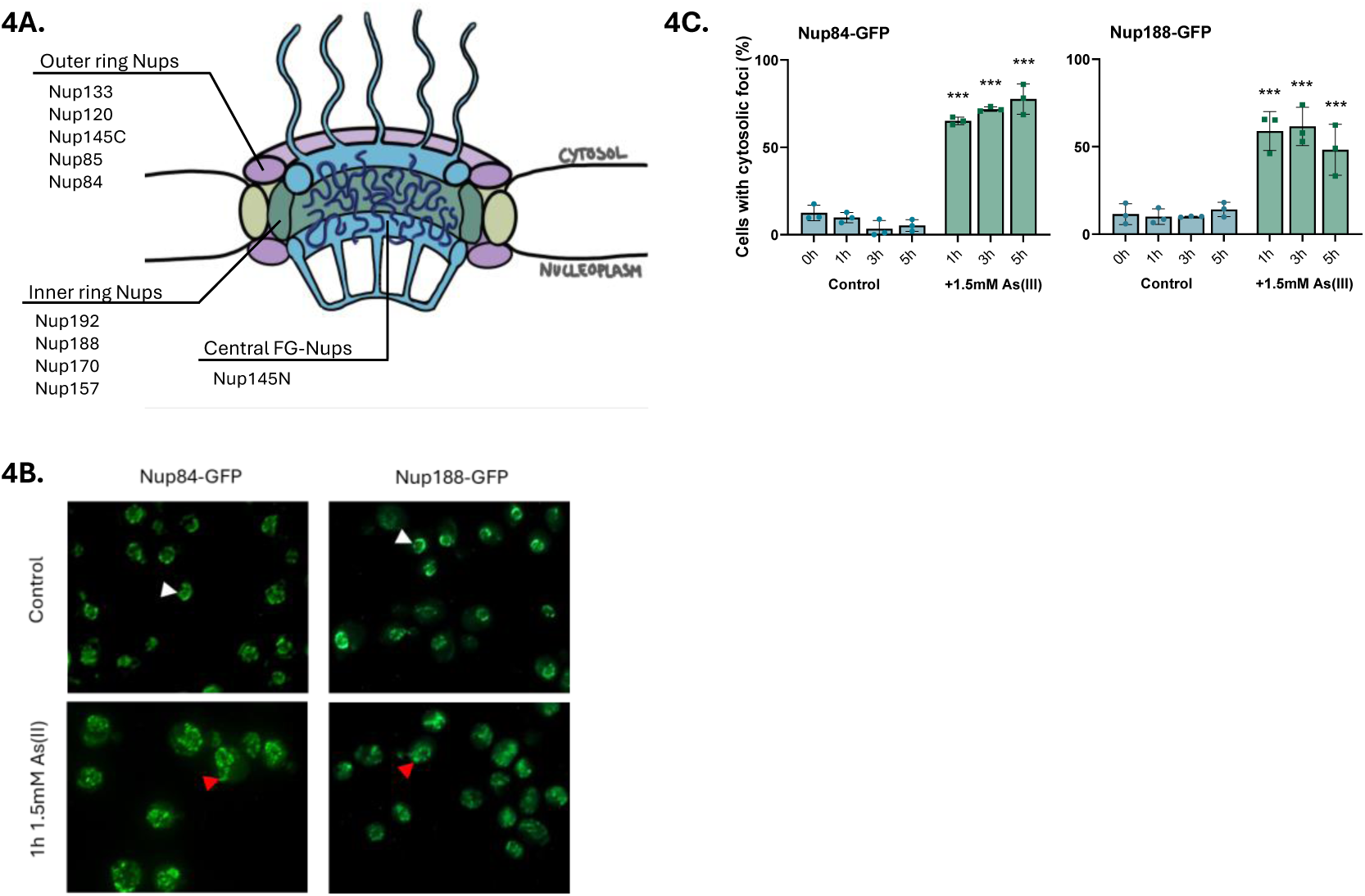
As(III) affects Nup localization in cells. **4A.** Illustration of the yeast NPC with the arsenic-binding Nups and their respective subcomplex. **4B**. Localization of GFP-tagged Nup84 and Nup188. In unexposed (control) cells the proteins are located around the NE (white arrowheads). After 1 h of As(III) exposure (1.5 mM), the proteins are visible as cytosolic foci, often with the fluorescence signal extending from the NE into the cytosol (red arrowheads). The images shown are representative of three biological repeats of 100 cells each. **4C**. Nup84–GFP and Nup188-GFP distribution was scored by fluorescence microscopy and quantified as in Figure 3B. The bars represent the mean ± SD of three independent biological repeats of a total of 300 cells. *P*-values are according to * > 0.05, ** > 0.01, *** > 0.001.

We next performed immuno-electron microscopy (EM) on untreated and As(III) exposed yeast cells using a primary antibody that detects Nups in conjunction with a secondary 10 nm gold label to simultaneously observe NE morphology, Nup localization, NPC morphology and number, and protein aggregates visible as electron-dense content (EDC) within cells (Panagaki et al., 2021; Schneider et al., 2024). While we did not detect any abnormalities in NPC morphology in As(III)-exposed cells (Fig. 5A), the exposed cells had fewer visible NPCs per cell section compared to unexposed cells (Fig. 5B). The lower number of NPCs is probably not a result of reduced Nup levels during As(III) stress as Nup84-GFP and Nup188-GFP levels remained largely unchanged during exposure (Fig. S4B). Instead, a substantial fraction of Nup immunolabelling was associated with EDCs in As(III)-exposed cells (Fig. 5C), suggesting that Nups may aggregate. These aggregates were not membrane enclosed, but often localized at sites of NE deformations in which the NE extended into the cytoplasm (Fig. 5D, panel I.) or where the inner and outer leaflets were separated with the outer leaflet extending into the cytoplasm (Fig. 5D, panel II.) forming an outer membrane bud. In sum, arsenic-binding to Nups may result in their mislocalization and aggregation, reducing the number of NPCs present on the NE.

**Figure 5.**
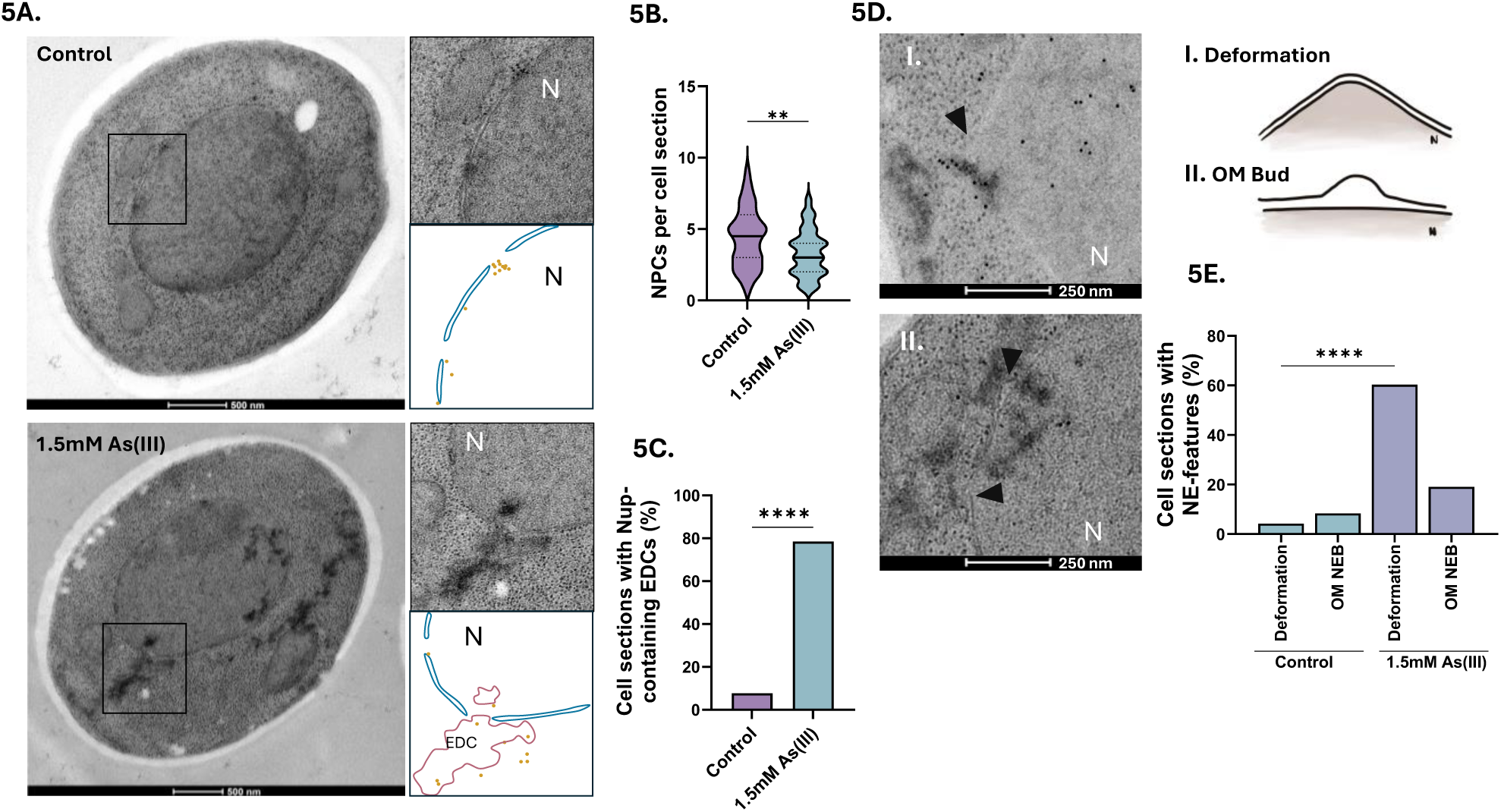
As(III) affects NE morphology and NPC numbers. **5A**. Representative electron micrographs of yeast cells before and after exposure to 1.5 mM As(III) for 1 h. The boxes correspond to areas showing NPCs that appear as holes in the NE bilayer. Nup localization was addressed with a gold-labelled anti-Nup antibody (Mab414). Protein aggregates appear as electron dense content (EDC) both in nucleus and in areas that are free of ribosomes in the cytosol (Panagaki et al., 2021). A model is drawn below each micrograph to visualize the locations of gold particles and EDCs. **5B-C**. Quantification of the number of NPCs per cell section (in **C**) and of the fraction of Nups associated with EDCs/protein aggregates (in **D**) in unexposed (control) and As(III)-exposed cells. Nups were detected using a gold-labelled anti-Nup antibody (Mab414). **5D**. Representative electron micrographs of yeast nuclei of cells exposed to 1.5 mM As(III) for 1 h showing NE deformations (arrowheads) in which the NE extends into the cytoplasm (panel I.) and where the inner and outer leaflets are separated with the outer leaflet extending into the cytoplasm (panel II.) forming an outer membrane bud (OM bud), often near EDCs/protein aggregates. A model is drawn to the right of each micrograph to visualize the two types of NE deformation. **5E**. Quantification of the fraction of NE deformations and outer membrane buds (OM bud) per cell section in unexposed (control) and As(III)-exposed cells. The number of cell sections assessed per condition was 62 for control and 74 for As(III) exposed cells. Significance was calculated using an unpaired t-test, and the *P*-values are according to: **> 0.01, ***> 0.001, and **** > 0.0001. N, nucleus; EDC, electron dense content, OM bud, outer membrane bud.

### Nuclear transport is disrupted during long-term As(III) stress

Having established that importins and Nups mislocalize and aggregate, we next addressed whether nuclear transport is affected in As(III)-exposed cells using established reporters (Barrientos et al., 2023; Rempel et al., 2019). We first monitored the localization of the Srp1/Kap95 substrate GFP-tcNLS (GFP with a tandem classical NLS) under the control of the galactose-inducible *GAL1* promoter. Expression of GFP-tcNLS was induced and localization of GFP-tcNLS determined by calculating the ratio of the fluorescence measured in the nucleus over the cytosol (N/C ratio) in the absence or presence of As(III) (Fig. 6A). Importantly, the N/C ratio significantly decreased in As(III)-exposed cells (Fig. 6B), suggesting that nuclear accumulation of GFP-tcNLS was inhibited. Similarly, the N/C ratios of two other reporters containing NLS sequences recognized respectively by Kap104 (Nab2NLS-GFP) and Kap121/Pse1 (Pho4NLS-GFP), also decreased in exposed cells. In contrast, the N/C ratio of GFP without an NLS remained largely unaffected during exposure (Fig. 6B). The total protein levels of Kap95, Srp1 and Kap121/Pse1 were not affected by As(III) (Fig. 3D). Thus, nuclear import of NLS-containing cargos (GFP-tcNLS, Nab2NLS-GFP, Pho4NLS-GFP) is disrupted in As(III) exposed cells, probably due to arsenic-binding to import receptors and Nups.

**Figure 6.**
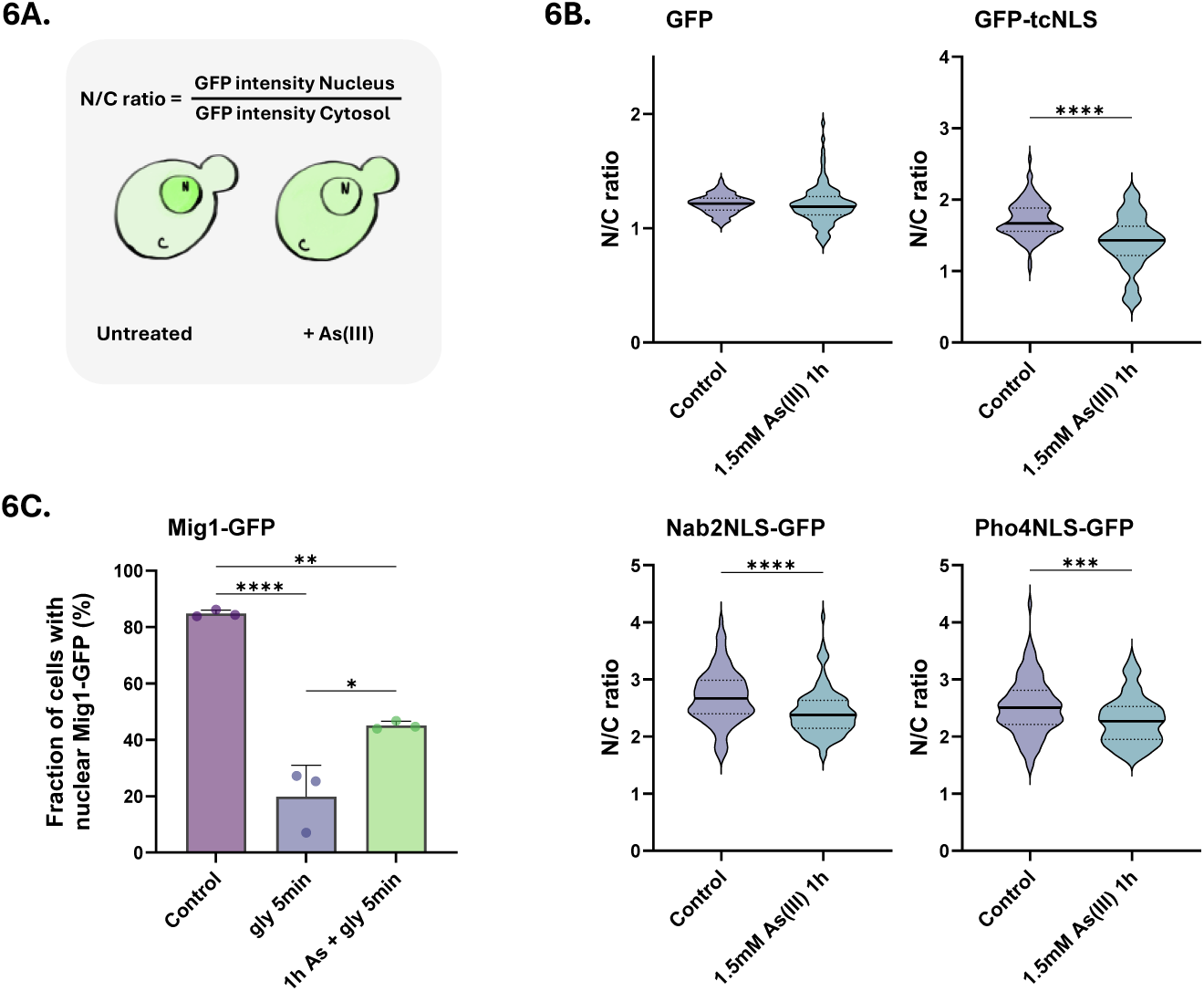
Nuclear transport is disrupted during As(III) stress. **6A**. Model of the assay. Yeast cells expressing GFP-tagged nuclear transport reporters were either left untreated or exposed to As(III). N/C ratios were calculated by measuring the mean fluorescence intensities in the nucleus (N) and the cytosol (C). **6B**. Nuclear protein import is inhibited by As(III). N/C ratios were determined in the absence (control) and presence of 1.5 mM As(III) for GFP-tcNLS (recognized by Kap95/Srp1), Nab2NLS-GFP (recognized by Kap104) and Pho4NLS-GFP (recognized by Kap121/Pse1). GFP without sorting sequence was included as a control. The graphs show the mean of three biological replicates of around 100 cells per condition measured. **6C**. Nuclear export is inhibited by As(III). Mig1-GFP localization was determined in the presence of glucose and 5 min after a shift to glycerol. Where indicated, cells were preincubated with 1.5 mM As(III) for 1 h before the shift to glycerol. Mig1-GFP distribution was scored by fluorescence microscopy and quantified as in Figure 3B. The bars represent the mean ± SD of three independent biological repeats of a total of 300 cells. Significance for 6B-C was calculated using an unpaired t-test, and *P*-values are according to: *> 0.05, **> 0.01, ***> 0.001, and **** > 0.0001.

To address if As(III) also affects nuclear export, we monitored the localization of the transcriptional repressor Mig1-GFP, which is nuclear in the presence of glucose but rapidly exits the nucleus via Msn5-dependent export when glucose is replaced by glycerol (DeVit and Johnston, 1999). Note that As(III) directly binds to Msn5 (Fig. 2B). As expected, shifting cells from glucose to glycerol resulted in a substantial drop of nuclear Mig1-GFP within 5 min (Fig. 6C). When cells were preincubated with As(III) for 1 h, glycerol-stimulated nuclear exit of Mig1-GFP was significantly inhibited (Fig. 6C). Thus, As(III) also disrupts nuclear export, possibly as a consequence of arsenic binding to export receptors.

Previous work has shown that yeast cells depend on transcriptional regulation of genes required for As(III) tolerance (Wysocki and Tamás, 2010). For instance, the transcription factors Yap1 and Msn2 accumulate in the nucleus during As(III) stress where they induce expression of defence genes (Hosiner et al., 2009; Thorsen et al., 2007; Wysocki et al., 2004) while the transcription factor Sfp1 exits the nucleus upon As(III) exposure which results in down-regulation of protein biosynthesis-related genes (Hosiner et al., 2009). Thus, cells rely on functional nucleocytoplasmic transport to mount an appropriate response to arsenic stress. Nuclear accumulation of chromosomally integrated Yap1-GFP and Msn2-GFP, as well as nuclear exit of Sfp1-GFP was efficient within 5-15 min of As(III) exposure (Fig. S6), consistent with their role as As(III)-responsive factors. These data suggest that nuclear protein import and export is not affected during short-term exposure, possibly because the arsenic that enters cells is first recognized by arsenic-specific sensing and signalling systems before sufficient arsenic accumulates to poison cellular functions. Alternatively, these stress-responsive transcription factors might cross the NE through As(III)-insensitive pathways. Collectively, our data suggest that nuclear transport is disrupted during long-term As(III) stress but remains functional in the initial phase of exposure.

Our chemical-genetic and genetic interaction data (Fig. S1) suggested that arsenic-binding to proteins mediating nucleocytoplasmic transport causes toxicity. In further support of this notion, cells with weakened (temperature-sensitive) alleles of Kap95 (*kap95*-L63A) and Kap121/Pse1 (*kap121*-Δ34) (Li et al., 2011) were As(III) sensitive (Fig. 7). Thus, disruption of nucleocytoplasmic transport may result in arsenic sensitivity.

**Figure 7.**
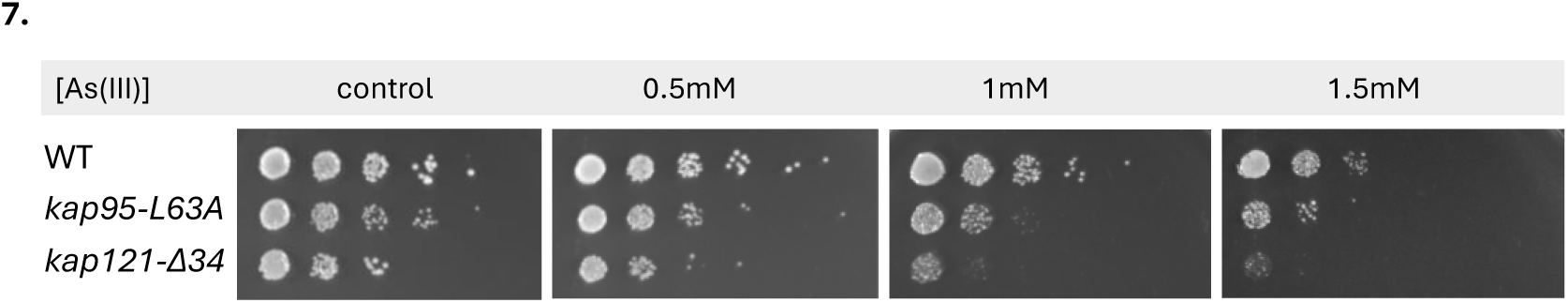
Cells defective in nuclear transport function are As(III) sensitive. Yeast cells that carry a weakened (temperature-sensitive) allele of Kap95 (*kap95-L63A*) or Kap121/Pse1 (*kap121-Δ34*) were grown to log phase, adjusted to the same OD, serially diluted, and plated onto YNB medium with 2% glucose as carbon source and the indicated As(III) concentrations. Plates were incubated at 30°C for 3 days. Images shown are representative of three biological replicates.

## DISCUSSION

This current study strongly implicates nucleocytoplasmic transport as a key target of arsenic toxicity. First, our proteome-wide approach identified 174 arsenic-binding proteins *in vivo*, of which proteins involved in nucleocytoplasmic transport were remarkably enriched. In fact, arsenic bound to most of the importins present in *S. cerevisiae* and we verified arsenic-binding to selected importins, exportins, and Nups. Second, we demonstrated that importins and Nups mislocalized and aggregated, and that the number of NPCs was reduced in As(III) exposed cells. Third, we provided evidence that nucleocytoplasmic transport is impaired during As(III) exposure and that cells with defective nuclear protein transport function are As(III) sensitive. Together, our data are consistent with a model in which arsenic-binding to nuclear transport factors leads to their mislocalization and aggregation, disrupting nucleocytoplasmic transport and causing As(III) sensitivity.

Previous *in vivo* proteome-wide studies using the As-biotin probe in various human cell lines typically identified 40-50 candidate arsenic-binding proteins (Dong et al., 2022; Yan et al., 2016; Zhang et al., 2015b; Zhang et al., 2007). Our current study yielded 174 proteins representing, to our knowledge, the largest set of *in vivo* arsenic-binding targets reported to date. Some proteins known to bind to arsenic were absent from our hit list, including the As(III)-sensing transcription factor Yap8 (Kumar et al., 2016). Low-abundance proteins are less represented in our dataset, and some true targets might be missed due to the stringent filtering criteria used. A study that applied As-biotin to a human proteome microarray identified 360 candidate arsenic-binding proteins (Zhang et al., 2015a). It is unclear whether proteins are properly folded on the microarray, which is an important aspect since non-native structures could result in cysteine residues being more easily accessible for arsenic-binding than in folded native proteins. Nevertheless, the authors demonstrated that one of the 360 candidate proteins, hexokinase 2, is a direct target of arsenic inhibition *in vivo* (Zhang et al., 2015a). Collectively, these studies demonstrate the great utility of the As-biotin probe for proteome-wide identification of binding and toxicity targets, and together provide a valuable resource for mechanistic studies of on the toxicity, pathology, and therapeutic effects of arsenicals.

While As-biotin is a valuable tool, it cannot discriminate between As(III)- and MAs(III)-binding proteins (Lee and Levin, 2019). By using *mtq2Δ* cells and pretreatments with As(III) and MAs(III) as blocking agents, we identified proteins that bound to either As(III) or MAs(III), or to both arsenicals (Table S2). For example, Kap121/Pse1 and Crm1 bound to both arsenicals whilst Kap95 primarily bound to MAs(III) (Fig. 2C). Interestingly, previous studies indicated that As(III) and MAs(III) can bind to distinct sets of cysteine thiols in target proteins thereby eliciting stress-specific responses (Lee and Levin, 2022). How specific cysteine residues distinguish between As(III) and MAs(III) is not known. While our study suggests that the majority of the 174 arsenic-binding proteins can bind to As(III) as well as MAs(III), the exact form of arsenic, the residues involved, and the physiological consequences of the binding remain to be defined.

The observations that As(III)-induced Kap95-GFP and Nup84-GFP foci formation were unaffected in the presence of CHX (Figs. 3B, S4), suggests that arsenic targets the native (folded) form of these proteins promoting their mislocalization and aggregation (Figs. 3, 4, 5). Previously, we showed that As(III) treatment of yeast cells resulted in the formation of Hsp104-GFP foci that could be prevented by CHX (Andersson et al., 2021; Hua et al., 2022; Ibstedt et al., 2014; Jacobson et al., 2012). Based on these and other findings, we concluded that As(III) primarily targets non-native proteins for misfolding and aggregation (Andersson et al., 2021; Hua et al., 2022; Ibstedt et al., 2014; Jacobson et al., 2012). It is important to note that there is a fundamental difference between Hsp104-GFP foci and Kap95-GFP foci that form in As(III)-exposed cells. Hsp104 is a disaggregase that associates with and reactivates aggregated proteins in *S. cerevisiae* (Glover and Lindquist, 1998) and Hsp104-GFP is a well-established marker for cytosolic protein aggregation (Jacobson et al., 2012; Kaganovich et al., 2008; Panagaki et al., 2021). Thus, As(III)-induced Hsp104-GFP foci formation is a consequence of global aggregation of cytosolic proteins to which Hsp104 associates while its own activity is unaffected (Hua et al., 2022; Jacobson et al., 2012). In contrast, Kap95-GFP foci represents aggregated (Fig. 3) and probably unfunctional Kap95 since nuclear import of the Kap95/Srp1 cargo GFP-tcNLS was impaired in As(III)-treated cells (Fig. 6B). Moreover, unlike Hsp104 and it cochaperones, karyopherins are unlikely to recognize misfolded proteins. Thus, native and functional Hsp104-GFP forms foci by associating with non-native proteins while Kap95-GFP (and probably also Nup84-GFP) most likely forms foci because it aggregates. Hence, arsenic can directly modify cysteines in non-native (Hua et al., 2022; Jacobson et al., 2012) as well as native (this work) proteins driving their unfolding and aggregation, with non-native proteins being particularly vulnerable (Tamás et al., 2014; Wysocki et al., 2023). This raises the question if some (or many) of the proteins isolated in the As-biotin experiments are being brought down indirectly in aggregates. However, the As-biotin pull-down experiments would probably not include aggregates because the beads are centrifuged at 1000 x *g*, which is not nearly high enough to sediment aggregates. This is also consistent with the relatively low level of overlap between the proteins identified as arsenic-binding and those identified by sedimentation of As(III)-induced aggregates (Fig. 1F).

It has been shown that cytosolic protein aggregates can impair nucleocytoplasmic transport by sequestering nuclear shuttle factors (Woerner et al., 2016). While As(III) induces global aggregation of cytosolic proteins in yeast (Jacobson et al., 2012), our data point to a more direct mechanism where arsenic impairs nucleocytoplasmic transport by binding to nuclear import and export factors. Arsenic may also affect nucleocytoplasmic transport by binding to Nups, which then leads to reduced number of NPCs in the NE (Fig. 5). Additionally, As(III) induces morphological abnormalities of the NE with deformations extending into the cytosol (Fig. 5). Interestingly, genetic perturbations that result in NE aberrations, such as NE protrusions that extend into the cytosol, are linked to loss-of-function mutations of NPC components (Thaller and Lusk, 2018) including several Nups identified in our study (*e.g*. Nup85, Nup188, Nup145). Thus, the observed NE aberration during As(III) stress may be a direct consequence of arsenic-binding to Nups, interfering with their function.

Several studies have linked arsenic exposure to an increased prevalence of neurodegenerative disorders such as Alzheimer’s and Parkinson’s disease (Rahman et al., 2021; Wysocki et al., 2023). These diseases are characterized by the pathological accumulation of protein aggregates (Soto and Pritzkow, 2018), and there is growing evidence that arsenic contributes to these diseases by impairing protein folding in cells (Wysocki et al., 2023). Interestingly, mislocalization of nucleocytoplasmic transport factors, such as Nups and importins, and the disruption of nucleocytoplasmic transport have been implicated in the pathology of neurodegenerative disorders (Hutten and Dormann, 2020; Kim and Taylor, 2017). Thus, the observed impairment of nucleocytoplasmic transport by arsenic, via direct binding, mislocalization, and aggregation of individual nuclear transport factors described in this current study, may be an additional mechanism by which this metalloid contributes to pathology.

In conclusion, our study provides novel insights into the molecular mechanisms by which arsenic disrupts cellular function, specifically its impact on nucleocytoplasmic transport. These findings have broad implications for understanding how environmental poisons affect cells and organisms and may serve as a basis for future research on the toxic and therapeutic effects of arsenicals.

## MATERIALS AND METHODS

### Yeast strains, plasmids, and culturing conditions

All yeast strains and plasmids used in this work are listed in Table S3. The *S. cerevisiae* strains are based on BY4741 (Brachmann et al., 1998), the deletion collection (Giaever et al., 2002), the collection of temperature-sensitive mutants of essential yeast genes (Li et al., 2011), and the GFP collection (Huh et al., 2003). The strains harbouring nuclear transport reporters have been described previously (Rempel et al., 2019).

Plasmids containing HA-tagged versions of Kap123, Sxm1, Nup84, and Nup188 were constructed via Gateway Recombination Cloning (Thermo Fisher Scientific, Waltham, MA, USA) in accordance with the manufacturer’s instructions. Gene sequences were amplified by PCR using genomic DNA as template, inserted into the donor plasmid pDONR221 (Thermo Fisher Scientific, Waltham, MA, USA) and then into the destination vector pAG426GPD-ccdB-HA (Addgene plasmid #14252; http://n2t.net/addgene:14252) (Alberti et al., 2007). All plasmids were verified by sequencing.

Cells were routinely grown at 30°C in minimal YNB (yeast nitrogen base) medium with 2% glucose as carbon source. Cells containing nuclear transport reporters were grown with 2% raffinose as carbon source and protein expression was induced with 0.5% galactose for 4 h. For growth assays on plates, cells were grown until log phase, the optical density (OD) at 600 nm adjusted to equal for all cultures, diluted in 10-fold steps, plated using a sterilized stamp, and incubated for 2–3 days at 30°C. Where indicated, the following chemicals were added: sodium arsenite (NaAsO_2_, S7400), cycloheximide (C7698) (both from Sigma-Aldrich, St. Louis, MO, USA), biotinyl p-aminophenyl arsenic acid (As-biotin, B394970; Toronto Research Chemicals, North York, Canada), and monomethylarsonous acid (CH_5_AsO_2_, M565100; LGC Standards, Wesel, Germany).

### Arsenic-binding assays

Proteome-wide identification of arsenic-binding proteins was performed with *mtq2Δ* cells grown to early log phase in YNB medium. For the last hour, the medium was switched to YEPD (Yeast Extract Peptone Dextrose) with 2% glucose and split into control and treatment cultures. The cultures were pretreated for 10 min with either As(III) or MAs(III) as blocking agents, followed by 10 min incubation with As-biotin at 30°C. The cultures were collected, pelleted, and frozen at -80°C or lysed directly. Cell lysis was performed by bead beating (1 min, 4°C) in IP buffer (1x TNT buffer (50 mM Tris-HCl, pH 7.5, 150 mM NaCl, 0.5% Triton, pH 7-7.5)), 1x protease inhibitor (Complete mini, EDTA-free; Roche), 1x phosphatase inhibitor (PhosSTOP Easypack; Roche), and 1 mM phenylmethylsulfonyl fluoride (PMSF). Streptavidin agarose beads (ThermoFisher, 20353) were prepared by a 1x TNT buffer wash and aliquoted to all lysates. After a 1 h incubation (4°C, rotating), pull-down was performed by centrifugation (1000x *g*, 1 min) and washing three times in 1x TNT. Proteins present in the pull-down were separated by SDS-PAGE electrophoresis followed by identification using microcapillary liquid chromatograph-tandem mass spectrometry (LC/MS/MS) at the Taplin Mass Spectrometry Facility at Harvard Medical School. To validate arsenic-binding to selected proteins, we used wild type cells carrying plasmids with HA-tagged versions of the corresponding genes or cells that harboured GFP-tagged versions of the genes in their genomes. After As-biotin pull-down and SDS-PAGE as described above, the proteins were visualized by Western blotting using anti-GFP rabbit IgG (1:8000, A11122; Invitrogen, Waltham, MA, USA) or anti-HA mouse IgG (1:1000, sc-7392; Santa Cruz Biotechnologies, Dallas, TX, USA) primary antibodies, and anti-rabbit IgG (1:3000, 84564) and anti-mouse IgG (1 : 5000, 84545) secondary antibodies (both from Invitrogen (Waltham, MA, USA)). A more extensive description of the Western blot protocol is provided in the section ‘Protein aggregate isolation and Western blotting’ below.

### Bioinformatics and protein structure analyses

Negative genetic interactors (including negative genetic, synthetic growth defect, synthetic lethality) of selected arsenic-binding hits were retrieved from the *Saccharomyces* genome database (SGD) (Wong et al., 2023) and compared to a compendium of 712 As(III) sensitive *S. cerevisiae* mutants that contains the genes identified at least once in four previous genome-wide phenotypic screens (Haugen et al., 2004; Pan et al., 2010; Thorsen et al., 2009; Zhou et al., 2009). The significance of the overlaps between datasets was calculated by the hyper-geometric test. All datasets used for comparisons are listed in Table S4. For protein structure analyses, protein sequences were retrieved from SGD. PDB files were retrieved from the AlphaFold protein structure database (Jumper et al., 2021) and visualized using UCSF ChimeraX version: 1.6.1 (Goddard et al., 2018).

### Fluorescence microscopy and image analyses

Cells expressing GFP-tagged proteins were grown until mid-log phase and either left untreated or exposed to As(III) and/or 0.2 mg/ml CHX added to the culture at the same time as the As(III). To induce nuclear export of Mig1-GFP, cells were first grown in glucose-containing medium and then shifted to medium containing 2% glycerol. Where indicated, cells were pretreated with 1.5 mM As(III) for 1 h. All samples were fixed in 3.7% formaldehyde (RT, 30 min) followed by two washes in 1x PBS. Nuclear staining was done by incubating fixed cells in ethanol (RT, 40 min), washing in 1x PBS, and resuspending in 4’,6-diamidino-2-phenylindole (DAPI) solution (D1306; Thermo Fisher Scientific). Fluorescent signals were detected using a Zeiss Axiovert 200 M (Carl Zeiss Microscopy, München, Germany) fluorescence microscope equipped with Plan-Apochromat 1.40 objectives and appropriate fluorescence light filter sets. Images were taken with a digital camera (AxioCamMR3). The ZEISS ZEN PRO software (Carl Zeiss Microscopy) was used to capture the images and the ImageJ-Fiji software (Schindelin et al., 2012) for quantifications.

The steady-state localization of nuclear transport reporters (GFP-tcNLS, Nab2NLS-GFP, Pho4NLS-GFP, GFP) was determined largely as described previously (Barrientos et al., 2023; Rempel et al., 2019). Cells were grown to mid-log phase in YNB medium containing 2% raffinose and then expression of the reporters was induced by the addition of 0.5% galactose for 4 h followed by the addition of 1.5 mM As(III) for 1 h. N/C ratios were quantified by measuring the mean fluorescence intensity in the nucleus and in the cytosol. The nucleus was outlined along the NE using Nup49-mCherry. Care was taken to exclude the vacuole when choosing a field in the cytosol. All measured values were corrected for background fluorescence and the ratio of nuclear versus cytosolic signal (N/C ratio) was calculated for three replicates and averaged.

### Protein aggregate isolation and Western blotting

Insoluble protein aggregates were isolated as described previously (Hua et al., 2022; Weids and Grant, 2014). Cells were grown to mid-log phase, unexposed or exposed to 1.5 mM As(III), collected by centrifugation, resuspended in lysis buffer (50 mM potassium phosphate buffer pH 7, 1 mM EDTA, 5% glycerol, 1 mM PMSF, EDTA-free protease inhibitor cocktail from Roche Diagnostics, Basel, Switzerland), and lysed with 2.5 mg/ml lyticase (Sigma-Aldrich: 30 min, 30°C). Cells were disrupted using sonication on ice (Sonifier 150, Branson Ultrasonics, Danbury, CT, USA; 8x 5 s pulses, 50% amplitude) and the total lysates collected by centrifugation. Protein concentrations in the lysates were adjusted to equal for all samples. Aggregated proteins were isolated by centrifugation of the protein lysates, resuspended in lysis buffer containing 20% NP40 twice, followed by washing and resuspension of the pellet in lysis buffer. The final resuspension was aided by sonication (2×5 sec, 50% amplitude on ice). 5X SDS loading buffer (4% SDS, 250 mM Tris Buffer pH 6.8, 16% β-mercaptoethanol, 30% glycerol, bromophenol blue) was added to all samples followed by boiling for 5 min at 95°C. Proteins were separated on a Criterion 4-20% stain free precast gel (Bio-Rad Laboratories, Hercules, CA, USA) and visualized using a ChemiDoc XRS+ system with UV-activation (Bio-Rad). For Western blot analysis, proteins were transferred to a PVDF membrane using TransBlot Turbo transfer system (semi-dry transfer, Bio-Rad). Membranes were blocked with 5% bovine serum albumin (BSA) in Tris-buffered saline (TBS) containing 0.05% Tween 20 (TBS-T) for 1 h at RT followed by over-night incubation at 4°C with an anti-GFP antibody (1:8000, A11122; Invitrogen, Waltham, MA, USA). Membranes were washed 3X with TBS-T, incubated for 2 h with anti-rabbit-IgG secondary antibody (1:5000, 84546; Invitrogen, Waltham, MA, USA) and washed with TBS-T followed by signal detection using the ChemiDoc system.

### Immuno-electron microscopy

Immuno-EM was performed largely as described previously (Panagaki et al., 2021). Cells were grown to early log phase and either left untreated or exposed to 1.5 mM As(III) for 1 h. Samples were high-pressure frozen (Wohlwend HPF Compact 3, Sennwald, Switzerland) followed by freeze substitution in 2% uranyl acetate (UA) dissolved in acetone, and embedded into HM20 lowicryl resin (Polysciences) that was UV polymerised at -50°C. The resin was sectioned in 70 nm thin sections and placed on mesh grids. The sections were fixed in 1% paraformaldehyde in PBS for 10 min and blocked with 0.1% fish skin gelatine and 0.8% BSA in PBS for 1 h. For detection of Nups, samples were incubated for 2 h with 1:120 dilution of the mouse monoclonal anti-NPC antibody Mab414 (Abcam ab24609) at 4°C, followed by incubations at room temperature with 1:150 dilution of rabbit anti-mouse immunoglobulin (Agilent/Dako, E0433) for 1 h and with 1:70 diluted 10 nm gold-conjugated protein A antibody (CMC UMC Utrecht, The Netherlands) for 30 min. Glutaraldehyde (2.5%) was applied to sections for 1 h, followed by contrast staining in 2% UA for 5 min and 1 min in Reynold’s lead citrate (Reynolds, 1963). Three washing steps (20 min, PBS) were carried out after incubations with each antibody. Images were acquired at 120 kV on a Tecnai T12 transmission electron microscope equipped with a Ceta CMOS 16M camera (Thermofischer scientific, Eindhoven, the Netherlands). Quantifications were made using IMOD (Kremer et al., 1996) and statistics with GraphPad Prism 10.

## Acknowledgements

We thank Liesbeth M. Veenhoff (University of Groningen) for providing yeast strains. pAG426GPD-ccdB-HA was a gift from Susan Lindquist (Addgene plasmid #14252; http://n2t.net/addgene:14252; RRID:Addgene_14252).

## Funding

This work was supported by grants from the National Institutes of Health (grant numbers R01GM138413 and R01GM48533) to D.E.L., Cancerfonden (grant number 211-865) and the Swedish Research Council (grant number 2019-4004) to J.L.H., the foundations Bengt Lundqvist minne och Sven och Dagmar Saléns Stiftelse to E.L., and the Swedish Research Council (grant number 2018-03577) and Olle Engkvists Stiftelse (grant number 216-0452) to M.J.T.

### Author contributions

D.E.L., J.L.H., and M.J.T. - conceptualization;

E.L., J.L., J.M., K.K., N.K., C.G., R.M. - investigation;

E.L., J.L., J.M., K.K., N.K., C.G., R.M., J.L.H., D.E.L., M.J.T. - formal analysis;

E.L. and M.J.T. – writing the original draft;

E.L., J.L., J.M., K.K., N.K., C.G., R.M., J.L.H., D.E.L., M.J.T. - review & editing;

E.L., D.E.L., J.L.H., and M.J.T. - funding acquisition.

## SUPPLEMENTARY MATERIAL

**Table S1.** All proteins detected by LC/MS/MS (data file).

**Table S2.** List of 174 candidate arsenic-binding proteins.

**Table S3**. List of strains and plasmids used.

**Table S4**. Datasets used for comparisons.

**Figure S1.**
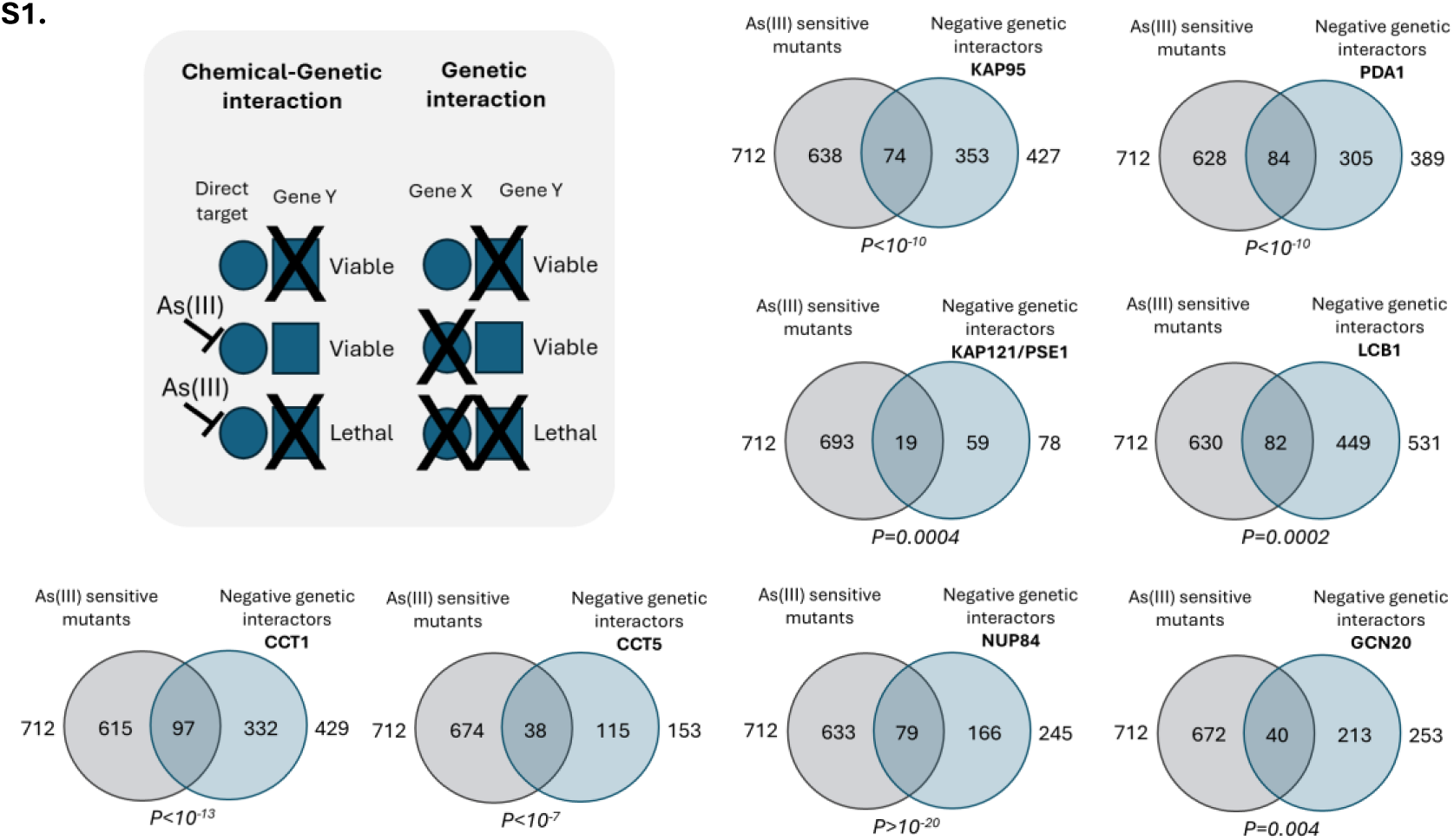
Integration of chemical-genetic and genetic interaction data to identify *bona fide* arsenic toxicity targets. Negative genetic interactors (including negative genetic, synthetic growth defect, synthetic lethality) of selected arsenic-binding hits were retrieved from SGD (Wong et al., 2023) and compared to a compendium of 712 As(III) sensitive *S. cerevisiae* mutants that contains the genes identified at least once in four genome-wide phenotypic screens (Haugen et al., 2004; Pan et al., 2010; Thorsen et al., 2009; Zhou et al., 2009). The significance of the overlaps between the datasets (negative genetic interactor sets and As(III) sensitive set) was calculated by the hyper-geometric test and the corresponding *P*-values are indicated.

**Figure S2.**
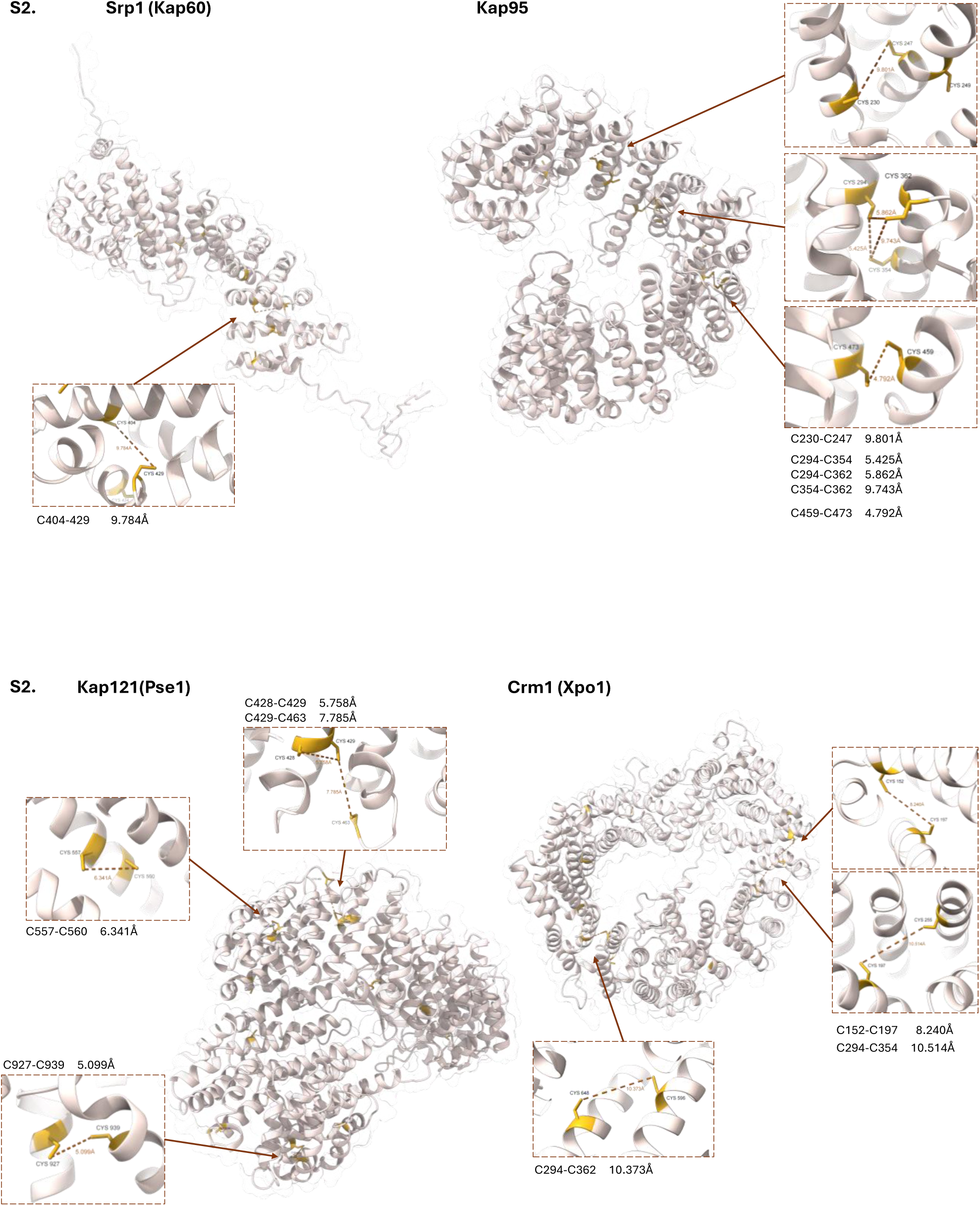
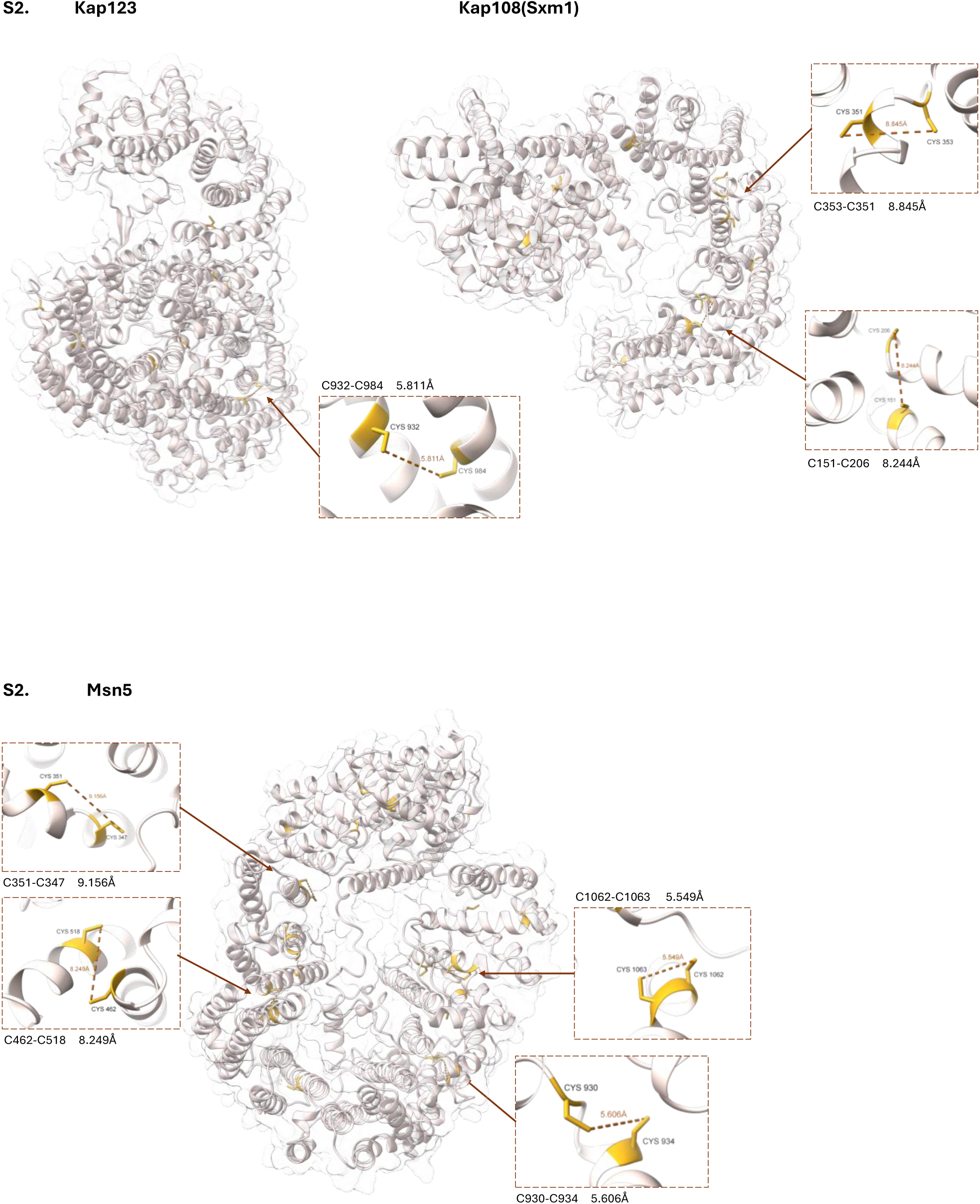
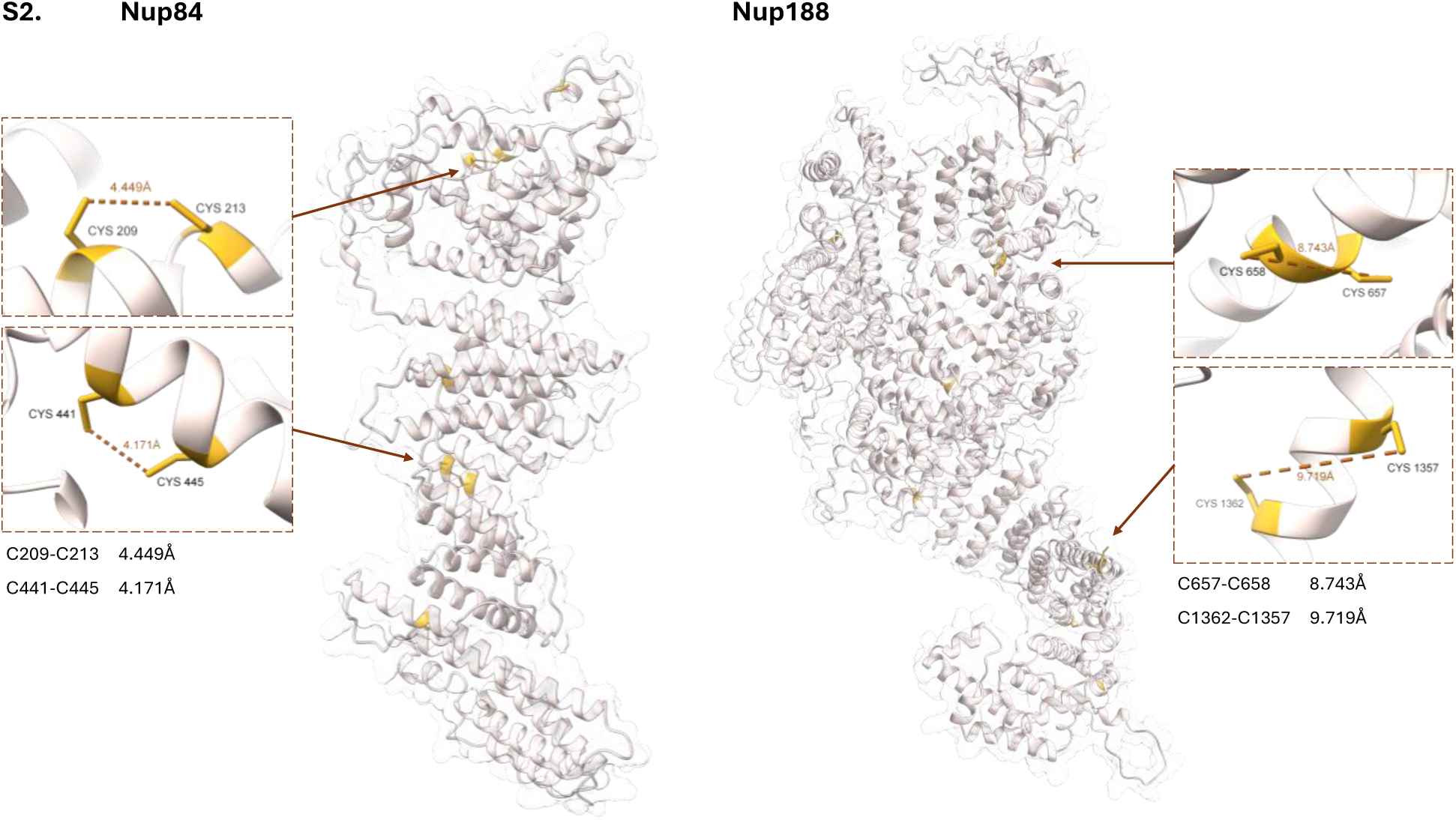
AlphaFold structure predictions and cysteine mapping for Srp1, Kap95, Kap121/Pse1, Crm1, Kap123, Sxm1/Kap108, Msn5, Nup84, and Nup188. Distances between pairs of adjacent or proximal cysteines are indicated. The structure predictions are based on experimental crystal structure data for all proteins except for Kap123, Kap108/Sxm1 and Msn5.

**Figure S3.**
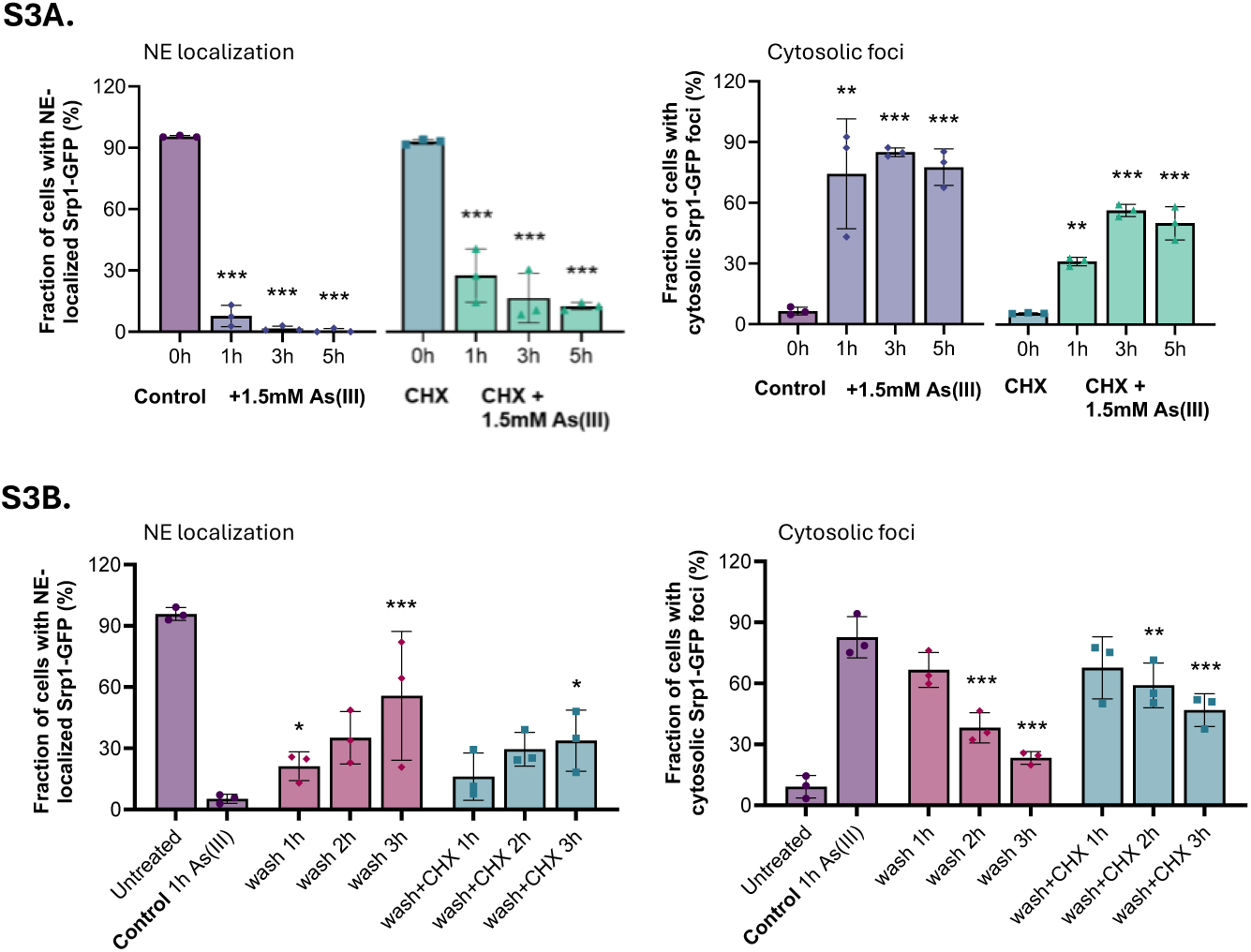
Mislocalization of Srp1 in As(III)-exposed cells. **S3A**. Quantification of Srp1-GFP nuclear envelope (NE) localization (left panel) and foci formation (right panel) in the presence and absence of 1.5 mM As(III) and/or 0.2 mg/ml cycloheximide (CHX). Srp1–GFP distribution was scored by fluorescence microscopy and quantified by visual inspection. The bars represent the mean ± SD of three independent biological repeats of a total of 300 cells. Significance was calculated using un-paired two-tailed student’s t-test with either the untreated control (for just As(III)-exposure) or CHX (for CHX+As(III) treated cells) as the comparison, and *P*-values are according to: ** > 0.01, *** > 0.001. **S3B**. Cells were exposed to 1.5 mM As(III) for 1 h, then washed twice and resuspended in medium without As(III) in the presence or absence of 0.2 mg/ml CHX. Srp1–GFP distribution was scored by fluorescence microscopy and quantified as in S3A. Significance was calculated using un-paired two-tailed student’s t-test of three independent biological replicates, with 1 h As(III)-exposed cells as the control sample. *P*-values according to * > 0.05, ** > 0.01, *** > 0.001.

**Figure S4.**
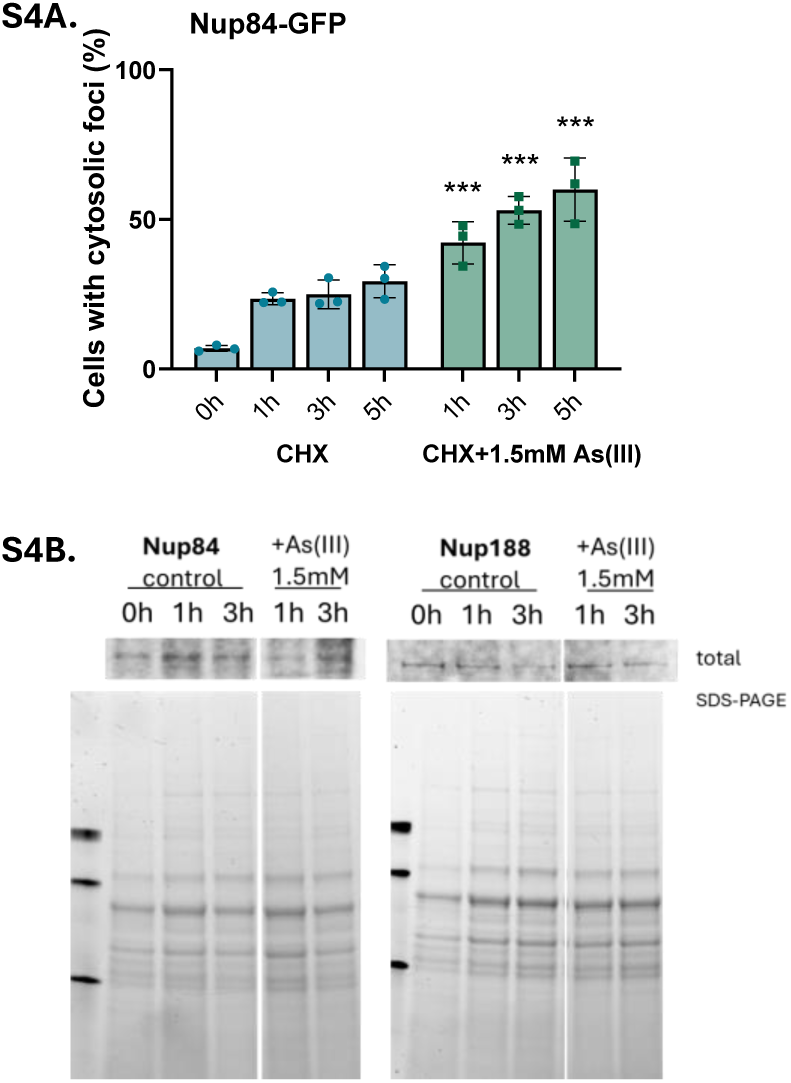
Nup localization and levels during As(III) stress. **S4A.** Cycloheximide (CHX) does not prevent Nup84-GFP mislocalization in As(III)-stressed cells. Quantification of Nup84-GFP foci formation in the presence or absence of 1.5 mM As(III) and 0.2 mg/ml CHX. Foci formation was detected with fluorescence microscopy and quantified by visual inspection. The bars represent the mean ± SD of three independent biological repeats of a total of 300 cells. Significance was calculated using un-paired two-tailed student’s t-test with CHX-treated cells as the comparison, and P-values are according to: ** > 0.01, *** > 0.001. **S4B.** Nup84-GFP and Nup188-GFP protein levels are largely unaffected by As(III). Western blot of the total lysate from cells expressing Nup84-GFP or Nup188-GFP in the absence (control) and presence of As(III). The lower panel shows the corresponding SDS-PAGE gels as loading controls. The images shown are representative of at least two biological repeats.

**Figure S5.**
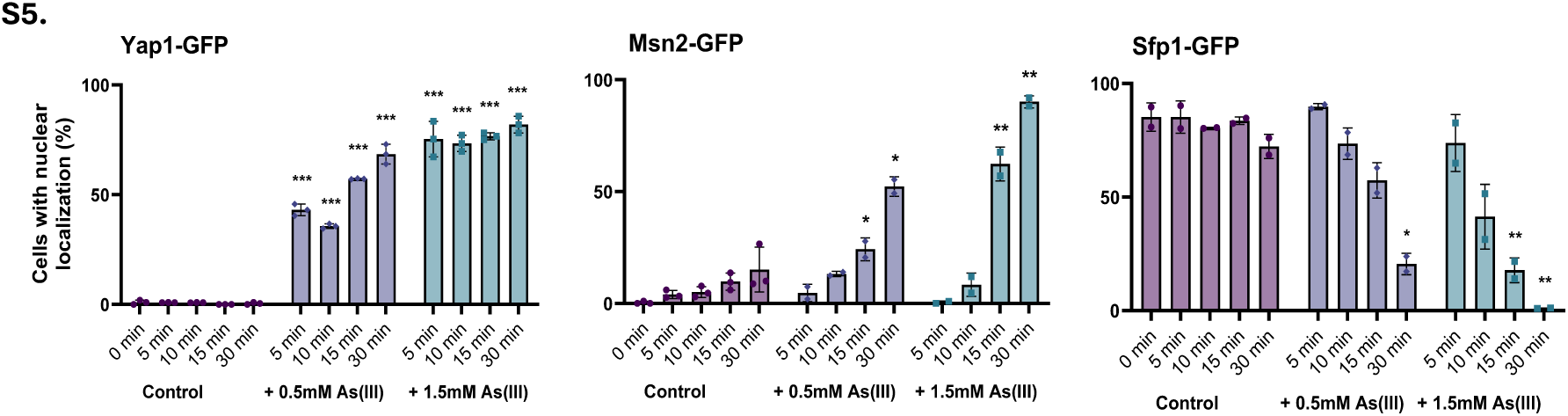
Nuclear transport is unaffected during short-term As(III) exposure. Cells expressing GFP-tagged versions of the transcription factors Yap1, Msn2, and Sfp1 were either left untreated (control) or exposed to the indicated concentrations of As(III), and their localization determined by fluorescence microscopy. Quantification was done by visual inspection and the bars represent the mean ± SD of three independent biological repeats of a total of 300 cells. Significance was calculated using un-paired two-tailed student’s t-test with the untreated control at the respective time point as the comparison, and *P*-values are according to: *> 0.05, ** > 0.01, *** > 0.001.

## REFERENCES

1. Aitchison, J.D., and M.P. Rout. 2012. The yeast nuclear pore complex and transport through it. Genetics. 190:855–883.

2. Alberti, S., A.D. Gitler, and S. Lindquist. 2007. A suite of Gateway cloning vectors for high-throughput genetic analysis in *Saccharomyces cerevisiae*. Yeast. 24:913–919.

3. Alberti, S., and A.A. Hyman. 2021. Biomolecular condensates at the nexus of cellular stress, protein aggregation disease and ageing. Nat Rev Mol Cell Biol. 22:196–213.

4. Andersson, S., A. Romero, J.I. Rodrigues, S. Hua, X. Hao, T. Jacobson, V. Karl, N. Becker, A. Ashouri, S. Rauch, T. Nyström, B. Liu, and M.J. Tamás. 2021. Genome-wide imaging screen uncovers molecular determinants of arsenite-induced protein aggregation and toxicity. J Cell Sci. 134:jcs258338.

5. Barrientos, E.C.R., T.A. Otto, S.N. Mouton, A. Steen, and L.M. Veenhoff. 2023. A survey of the specificity and mechanism of 1,6 hexanediol-induced disruption of nuclear transport. Nucleus. 14:2240139.

6. Bergquist, E.R., R.J. Fischer, K.D. Sugden, and B.D. Martin. 2009. Inhibition by methylated organo-arsenicals of the respiratory 2-oxo-acid dehydrogenases. J Organomet Chem. 694:973–980.

7. Brachmann, C.B., A. Davies, G.J. Cost, E. Caputo, J. Li, P. Hieter, and J.D. Boeke. 1998. Designer deletion strains derived from *Saccharomyces cerevisiae* S288C: a useful set of strains and plasmids for PCR-mediated gene disruption and other applications. Yeast. 14:115–132.

8. Chen, Q.Y., and M. Costa. 2021. Arsenic: A Global Environmental Challenge. Annu Rev Pharmacol. 61:47–63.

9. Chen, Q.Y., T. DesMarais, and M. Costa. 2019. Metals and Mechanisms of Carcinogenesis. Annu Rev Pharmacol Toxicol. 59:537–554.

10. DeVit, M.J., and M. Johnston. 1999. The nuclear exportin Msn5 is required for nuclear export of the Mig1 glucose repressor of *Saccharomyces cerevisiae*. Curr Biol. 9:1231–1241.

11. Dong, X., P. Wang, and Y. Wang. 2022. Chemoproteomic Approach for the Quantitative Identification of Arsenic-Binding Proteins. Chem Res Toxicol. 35:2145–2151.

12. Giaever, G., A.M. Chu, L. Ni, C. Connelly, L. Riles, S. Veronneau, S. Dow, A. Lucau-Danila, K. Anderson, B. Andre, A.P. Arkin, A. Astromoff, M. El-Bakkoury, R. Bangham, R. Benito, S. Brachat, S. Campanaro, M. Curtiss, K. Davis, A. Deutschbauer, K.D. Entian, P. Flaherty, F. Foury, D.J. Garfinkel, M. Gerstein, D. Gotte, U. Guldener, J.H. Hegemann, S. Hempel, Z. Herman, D.F. Jaramillo, D.E. Kelly, S.L. Kelly, P. Kotter, D. LaBonte, D.C. Lamb, N. Lan, H. Liang, H. Liao, L. Liu, C. Luo, M. Lussier, R. Mao, P. Menard, S.L. Ooi, J.L. Revuelta, C.J. Roberts, M. Rose, P. Ross-Macdonald, B. Scherens, G. Schimmack, B. Shafer, D.D. Shoemaker, S. Sookhai-Mahadeo, R.K. Storms, J.N. Strathern, G. Valle, M. Voet, G. Volckaert, C.Y. Wang, T.R. Ward, J. Wilhelmy, E.A. Winzeler, Y. Yang, G. Yen, E. Youngman, K. Yu, H. Bussey, J.D. Boeke, M. Snyder, P. Philippsen, R.W. Davis, and M. Johnston. 2002. Functional profiling of the *Saccharomyces cerevisiae* genome. Nature. 418:387–391.

13. Giaever, G., D.D. Shoemaker, T.W. Jones, H. Liang, E.A. Winzeler, A. Astromoff, and R.W. Davis. 1999. Genomic profiling of drug sensitivities via induced haploinsufficiency. Nat Genet. 21:278–283.

14. Glover, J.R., and S. Lindquist. 1998. Hsp104, Hsp70, and Hsp40: a novel chaperone system that rescues previously aggregated proteins. Cell. 94:73–82.

15. Goddard, T.D., C.C. Huang, E.C. Meng, E.F. Pettersen, G.S. Couch, J.H. Morris, and T.E. Ferrin. 2018. UCSF ChimeraX: Meeting modern challenges in visualization and analysis. Protein Sci. 27:14–25.

16. Guerra-Moreno, A., M.A. Prado, J. Ang, H.M. Schnell, Y. Micoogullari, J.A. Paulo, D. Finley, S.P. Gygi, and J. Hanna. 2019. Thiol-based direct threat sensing by the stress-activated protein kinase Hog1. Sci Signal. 12:eaaw4956.

17. Haugen, A.C., R. Kelley, J.B. Collins, C.J. Tucker, C. Deng, C.A. Afshari, J.M. Brown, T. Ideker, and B. Van Houten. 2004. Integrating phenotypic and expression profiles to map arsenic-response networks. Genome Biol. 5:R95.

18. Ho, B., A. Baryshnikova, and G.W. Brown. 2018. Unification of Protein Abundance Datasets Yields a Quantitative *Saccharomyces cerevisiae* Proteome. Cell Syst. 6:192–205 e193.

19. Hosiner, D., H. Lempiainen, W. Reiter, J. Urban, R. Loewith, G. Ammerer, R. Schweyen, D. Shore, and C. Schuller. 2009. Arsenic toxicity to *Saccharomyces cerevisiae* is a consequence of inhibition of the TORC1 kinase combined with a chronic stress response. Mol Biol Cell. 20:1048–1057.

20. Hua, S., A. Klosowska, J.I. Rodrigues, G. Petelski, L.A. Esquembre, E. Lorentzon, L.F. Olsen, K. Liberek, and M.J. Tamás. 2022. Differential contributions of the proteasome, autophagy, and chaperones to the clearance of arsenite-induced protein aggregates in yeast. J Biol Chem. 298:102680.

21. Huh, W.K., J.V. Falvo, L.C. Gerke, A.S. Carroll, R.W. Howson, J.S. Weissman, and E.K. O’Shea. 2003. Global analysis of protein localization in budding yeast. Nature. 425:686–691.

22. Hutten, S., and D. Dormann. 2020. Nucleocytoplasmic transport defects in neurodegeneration - Cause or consequence? Semin Cell Dev Biol. 99:151–162.

23. Ibstedt, S., T.C. Sideri, C.M. Grant, and M.J. Tamás. 2014. Global analysis of protein aggregation in yeast during physiological conditions and arsenite stress. Biology open. 3:913–923.

24. Jacobson, T., C. Navarrete, S.K. Sharma, T.C. Sideri, S. Ibstedt, S. Priya, C.M. Grant, P. Christen, P. Goloubinoff, and M.J. Tamás. 2012. Arsenite interferes with protein folding and triggers formation of protein aggregates in yeast. J Cell Sci. 125:5073–5083.

25. Jumper, J., R. Evans, A. Pritzel, T. Green, M. Figurnov, O. Ronneberger, K. Tunyasuvunakool, R. Bates, A. Zidek, A. Potapenko, A. Bridgland, C. Meyer, S.A.A. Kohl, A.J. Ballard, A. Cowie, B. Romera-Paredes, S. Nikolov, R. Jain, J. Adler, T. Back, S. Petersen, D. Reiman, E. Clancy, M. Zielinski, M. Steinegger, M. Pacholska, T. Berghammer, S. Bodenstein, D. Silver, O. Vinyals, A.W. Senior, K. Kavukcuoglu, P. Kohli, and D. Hassabis. 2021. Highly accurate protein structure prediction with AlphaFold. Nature. 596:583–589.

26. Kaganovich, D., R. Kopito, and J. Frydman. 2008. Misfolded proteins partition between two distinct quality control compartments. Nature. 454:1088–1095.

27. Kim, H.J., and J.P. Taylor. 2017. Lost in Transportation: Nucleocytoplasmic Transport Defects in ALS and Other Neurodegenerative Diseases. Neuron. 96:285–297.

28. Kitchin, K.T., and K. Wallace. 2008. The role of protein binding of trivalent arsenicals in arsenic carcinogenesis and toxicity. J Inorg Biochem. 102:532–539.

29. Kremer, J.R., D.N. Mastronarde, and J.R. McIntosh. 1996. Computer visualization of three-dimensional image data using IMOD. J Struct Biol. 116:71–76.

30. Kumar, N.V., J. Yang, J.K. Pillai, S. Rawat, C. Solano, A. Kumar, M. Grotli, T.L. Stemmler, B.P. Rosen, and M.J. Tamás. 2016. Arsenic directly binds to and activates the yeast AP-1-like transcription factor Yap8. Mol Cell Biol. 36:913–922.

31. Lallemand-Breitenbach, V., M. Jeanne, S. Benhenda, R. Nasr, M. Lei, L. Peres, J. Zhou, J. Zhu, B. Raught, and H. de The. 2008. Arsenic degrades PML or PML-RARalpha through a SUMO-triggered RNF4/ubiquitin-mediated pathway. Nat Cell Biol. 10:547–555.

32. Lee, J., and D.E. Levin. 2018. Intracellular mechanism by which arsenite activates the yeast stress MAPK Hog1. Mol Biol Cell. 29:1904–1915.

33. Lee, J., and D.E. Levin. 2019. Methylated metabolite of arsenite blocks glycerol production in yeast by inhibition of glycerol-3-phosphate dehydrogenase. Mol Biol Cell. 30:2134–2140.

34. Lee, J., and D.E. Levin. 2022. Differential metabolism of arsenicals regulates Fps1-mediated arsenite transport. J Cell Biol. 221:e202109034.

35. Li, Z., F.J. Vizeacoumar, S. Bahr, J. Li, J. Warringer, F.S. Vizeacoumar, R. Min, B. Vandersluis, J. Bellay, M. Devit, J.A. Fleming, A. Stephens, J. Haase, Z.Y. Lin, A. Baryshnikova, H. Lu, Z. Yan, K. Jin, S. Barker, A. Datti, G. Giaever, C. Nislow, C. Bulawa, C.L. Myers, M. Costanzo, A.C. Gingras, Z. Zhang, A. Blomberg, K. Bloom, B. Andrews, and C. Boone. 2011. Systematic exploration of essential yeast gene function with temperature-sensitive mutants. Nat Biotechnol. 29:361–367.

36. Liu, J., B. Chen, R. Zhang, Y. Li, R. Chen, S. Zhu, S. Wen, and T. Luan. 2023. Recent progress in analytical strategies of arsenic-binding proteomes in living systems. Anal Bioanal Chem. 415:6915–6929.

37. Lum, P.Y., C.D. Armour, S.B. Stepaniants, G. Cavet, M.K. Wolf, J.S. Butler, J.C. Hinshaw, P. Garnier, G.D. Prestwich, A. Leonardson, P. Garrett-Engele, C.M. Rush, M. Bard, G. Schimmack, J.W. Phillips, C.J. Roberts, and D.D. Shoemaker. 2004. Discovering modes of action for therapeutic compounds using a genome-wide screen of yeast heterozygotes. Cell. 116:121–137.

38. Marino, S.M., Y. Li, D.E. Fomenko, N. Agisheva, R.L. Cerny, and V.N. Gladyshev. 2010. Characterization of surface-exposed reactive cysteine residues in *Saccharomyces cerevisiae*. Biochemistry. 49:7709–7721.

39. Pan, X., S. Reissman, N.R. Douglas, Z. Huang, D.S. Yuan, X. Wang, J.M. McCaffery, J. Frydman, and J.D. Boeke. 2010. Trivalent arsenic inhibits the functions of chaperonin complex. Genetics. 186:725–734.

40. Panagaki, D., J.T. Croft, K. Keuenhof, L. Larsson Berglund, S. Andersson, V. Köhler, S. Büttner, M.J. Tamás, T. Nyström, R. Neutze, and J.L. Höög. 2021. Nuclear envelope budding is a response to cellular stress. Proc Natl Acad Sci U S A. 118:e2020997118.

41. Parsons, A.B., R.L. Brost, H.M. Ding, Z.J. Li, C.Y. Zhang, B. Sheikh, G.W. Brown, P.M. Kane, T.R. Hughes, and C. Boone. 2004. Integration of chemical-genetic and genetic interaction data links bioactive compounds to cellular target pathways. Nature Biotechnology. 22:62–69.

42. Paul, N.P., A.E. Galvan, K. Yoshinaga-Sakurai, B.P. Rosen, and M. Yoshinaga. 2023. Arsenic in medicine: past, present and future. Biometals. 36:283–301.

43. Peters, R.A., H.M. Sinclair, and R.H. Thompson. 1946. An analysis of the inhibition of pyruvate oxidation by arsenicals in relation to the enzyme theory of vesication. Biochem J. 40:516–524.

44. Rahman, M.A., M.A. Hannan, M.J. Uddin, M.S. Rahman, M.M. Rashid, and B. Kim. 2021. Exposure to Environmental Arsenic and Emerging Risk of Alzheimer’s Disease: Perspective Mechanisms, Management Strategy, and Future Directions. Toxics. 9:188.

45. Ramadan, D., P.C. Rancy, R.P. Nagarkar, J.P. Schneider, and C. Thorpe. 2009. Arsenic(III) species inhibit oxidative protein folding *in vitro*. Biochemistry. 48:424–432.

46. Rempel, I.L., M.M. Crane, D.J. Thaller, A. Mishra, D.P. Jansen, G. Janssens, P. Popken, A. Aksit, M. Kaeberlein, E. van der Giessen, A. Steen, P.R. Onck, C.P. Lusk, and L.M. Veenhoff. 2019. Age-dependent deterioration of nuclear pore assembly in mitotic cells decreases transport dynamics. eLife. 8:e48186.

47. Reynolds, E.S. 1963. The use of lead citrate at high pH as an electron-opaque stain in electron microscopy. J Cell Biol. 17:208–212.

48. Sapra, A., D. Ramadan, and C. Thorpe. 2015. Multivalency in the inhibition of oxidative protein folding by arsenic(III) species. Biochemistry. 54:612–621.

49. Schindelin, J., I. Arganda-Carreras, E. Frise, V. Kaynig, M. Longair, T. Pietzsch, S. Preibisch, C. Rueden, S. Saalfeld, B. Schmid, J.Y. Tinevez, D.J. White, V. Hartenstein, K. Eliceiri, P. Tomancak, and A. Cardona. 2012. Fiji: an open-source platform for biological-image analysis. Nat Methods. 9:676–682.

50. Schneider, K.L., X. Hao, K.S. Keuenhof, L.L. Berglund, A. Fischbach, D. Ahmadpour, S. Chawla, P. Gomez, J.L. Hoog, P.O. Widlund, and T. Nystrom. 2024. Elimination of virus-like particles reduces protein aggregation and extends replicative lifespan in *Saccharomyces cerevisiae*. Proc Natl Acad Sci U S A. 121:e2313538121.

51. Shen, S., X.F. Li, W.R. Cullen, M. Weinfeld, and X.C. Le. 2013. Arsenic Binding to Proteins. Chem Rev. 113:7769–7792.

52. Shi, W., J. Wu, and B.P. Rosen. 1994. Identification of a putative metal binding site in a new family of metalloregulatory proteins. J Biol Chem. 269:19826–19829.

53. Soto, C., and S. Pritzkow. 2018. Protein misfolding, aggregation, and conformational strains in neurodegenerative diseases. Nat Neurosci. 21:1332–1340.

54. Tamás, M.J., K.S. Sharma, S. Ibstedt, T. Jacobson, and P. Christen. 2014. Heavy metals and metalloids as a cause for protein misfolding and aggregation. Biomolecules. 4:252–267.

55. Thaller, D.J., and C.P. Lusk. 2018. Fantastic nuclear envelope herniations and where to find them. Biochem Soc T. 46:877–889.

56. Thomas, D.J. 2021. Arsenic methylation - Lessons from three decades of research. Toxicology. 457:152800.

57. Thorsen, M., G. Lagniel, E. Kristiansson, C. Junot, O. Nerman, J. Labarre, and M.J. Tamás. 2007. Quantitative transcriptome, proteome, and sulfur metabolite profiling of the *Saccharomyces cerevisiae* response to arsenite. Physiol Genomics. 30:35–43.

58. Thorsen, M., G.G. Perrone, E. Kristiansson, M. Traini, T. Ye, I.W. Dawes, O. Nerman, and M.J. Tamás. 2009. Genetic basis of arsenite and cadmium tolerance in *Saccharomyces cerevisiae*. BMC Genomics. 10:105.

59. Vergara-Geronimo, C.A., A. Leon Del Rio, M. Rodriguez-Dorantes, P. Ostrosky-Wegman, and A.M. Salazar. 2021. Arsenic-protein interactions as a mechanism of arsenic toxicity. Toxicol Appl Pharmacol. 431:115738.

60. Wang, Y., E. Weisenhorn, C.W. MacDiarmid, C. Andreini, M. Bucci, J. Taggart, L. Banci, J. Russell, J.J. Coon, and D.J. Eide. 2018. The cellular economy of the *Saccharomyces cerevisiae* zinc proteome. Metallomics. 10:1755–1776.

61. Weids, A.J., and C.M. Grant. 2014. The yeast peroxiredoxin Tsa1 protects against protein-aggregate-induced oxidative stress. J Cell Sci. 127:1327–1335.

62. Wing, C.E., H.Y.J. Fung, and Y.M. Chook. 2022. Karyopherin-mediated nucleocytoplasmic transport. Nat Rev Mol Cell Biol. 23:307–328.

63. Woerner, A.C., F. Frottin, D. Hornburg, L.R. Feng, F. Meissner, M. Patra, J. Tatzelt, M. Mann, K.F. Winklhofer, F.U. Hartl, and M.S. Hipp. 2016. Cytoplasmic protein aggregates interfere with nucleocytoplasmic transport of protein and RNA. Science. 351:173–176.

64. Wong, E.D., S.R. Miyasato, S. Aleksander, K. Karra, R.S. Nash, M.S. Skrzypek, S. Weng, S.R. Engel, and J.M. Cherry. 2023. *Saccharomyces* genome database update: server architecture, pan-genome nomenclature, and external resources. Genetics. 224:iyac191.

65. Wysocki, R., P.K. Fortier, E. Maciaszczyk, M. Thorsen, A. Leduc, A. Odhagen, G. Owsianik, S. Ulaszewski, D. Ramotar, and M.J. Tamás. 2004. Transcriptional activation of metalloid tolerance genes in *Saccharomyces cerevisiae* requires the AP-1-like proteins Yap1p and Yap8p. Mol Biol Cell. 15:2049–2060.

66. Wysocki, R., J.I. Rodrigues, I. Litwin, and M.J. Tamás. 2023. Mechanisms of genotoxicity and proteotoxicity induced by the metalloids arsenic and antimony. Cell Mol Life Sci. 80:342.

67. Wysocki, R., and M.J. Tamás. 2010. How *Saccharomyces cerevisiae* copes with toxic metals and metalloids. FEMS Microbiol Rev. 34:925–951.

68. Yan, X., J. Li, Q. Liu, H. Peng, A. Popowich, Z. Wang, X.F. Li, and X.C. Le. 2016. p-Azidophenylarsenoxide: An Arsenical “Bait” for the In Situ Capture and Identification of Cellular Arsenic-Binding Proteins. Angew Chem Int Ed. 55:14051–14056.

69. Zhang, H.N., L. Yang, J.Y. Ling, D.M. Czajkowsky, J.F. Wang, X.W. Zhang, Y.M. Zhou, F. Ge, M.K. Yang, Q. Xiong, S.J. Guo, H.Y. Le, S.F. Wu, W. Yan, B. Liu, H. Zhu, Z. Chen, and S.C. Tao. 2015a. Systematic identification of arsenic-binding proteins reveals that hexokinase-2 is inhibited by arsenic. Proc Natl Acad Sci U S A. 112:15084–15089.

70. Zhang, T., H. Lu, W. Li, R. Hu, and Z. Chen. 2015b. Identification of Arsenic Direct-Binding Proteins in Acute Promyelocytic Leukaemia Cells. Int J Mol Sci. 16:26871–26879.

71. Zhang, X., F. Yang, J.Y. Shim, K.L. Kirk, D.E. Anderson, and X. Chen. 2007. Identification of arsenic-binding proteins in human breast cancer cells. Cancer Lett. 255:95–106.

72. Zhang, X.W., X.J. Yan, Z.R. Zhou, F.F. Yang, Z.Y. Wu, H.B. Sun, W.X. Liang, A.X. Song, V. Lallemand-Breitenbach, M. Jeanne, Q.Y. Zhang, H.Y. Yang, Q.H. Huang, G.B. Zhou, J.H. Tong, Y. Zhang, J.H. Wu, H.Y. Hu, H. de The, S.J. Chen, and Z. Chen. 2010. Arsenic trioxide controls the fate of the PML-RARalpha oncoprotein by directly binding PML. Science. 328:240–243.

73. Zhou, X., A. Arita, T.P. Ellen, X. Liu, J. Bai, J.P. Rooney, A.D. Kurtz, C.B. Klein, W. Dai, T.J. Begley, and M. Costa. 2009. A genome-wide screen in *Saccharomyces cerevisiae* reveals pathways affected by arsenic toxicity. Genomics. 94:294–307.

74. Zhou, X., X. Sun, K.L. Cooper, F. Wang, K.J. Liu, and L.G. Hudson. 2011. Arsenite interacts selectively with zinc finger proteins containing C3H1 or C4 motifs. J Biol Chem. 286:22855–22863.

75. Zhou, Y., B. Zhou, L. Pache, M. Chang, A.H. Khodabakhshi, O. Tanaseichuk, C. Benner, and S.K. Chanda. 2019. Metascape provides a biologist-oriented resource for the analysis of systems-level datasets. Nature Commun. 10:1523.

